# A novel role for the *CNTN6* locus in lumenization and radial glial cell fate determination during early human cortical development revealed in cerebral organoids

**DOI:** 10.1101/2025.10.09.681391

**Authors:** T.A. Shnaider, A.M. Yunusova, S.A Yakovleva, A.S. Knyazeva, I.E. Pristyazhnuk, P.S. Belokopytova, A.A. Khabarova, A.S. Ryzhkova, V.S. Fishman, T.V Nikitina, I.N. Lebedev, V.S. Tarabykin, A.V. Smirnov, O.L. Serov

## Abstract

Neurodevelopmental disorders are a class of heterogeneous diseases with a significant genetic contribution, including pathologies resulting from copy number variations (CNVs). Recent advancements in genetic diagnostic technologies have led to the identification of new genes associated with neurodevelopmental disorders through CNVs. One such gene is *CNTN6*, whose variants and CNVs are associated with intellectual disability and autism spectrum disorders. *Cntn6* encodes a neural cell adhesion molecule involved in rodents in axon and dendrite guidance, synapse formation, and oligodendrocyte differentiation, playing a critical role in brain development. However, in humans, the specific molecular and cellular pathogenetic mechanisms remain elusive. Using various techniques to model human brain development pathologies, such as somatic cell reprogramming, cerebral organoids, and genome editing, we established that the *CNTN6* locus is involved in the lumenization and cell identity of radial glial cells, as well as in regulating their proliferation. Furthermore, we found that the *CNTN6* locus is involved in the nuclear-cytoplasmic translocation of PAX6 protein, a key regulator of forebrain development. Molecular studies revealed that CNTN6 partially functions through the Notch signaling pathway during the early stages of human brain development. Our findings unveil a novel role of the *CNTN6* locus in the early stages of human cortical development.

## Introduction

Human brain development is a complex, multistage process with intricate and tight regulation. Alterations in this process can lead to the emergence of neurodevelopmental disorders – a diverse group of pathologies, including autism spectrum disorders (ASDs), intellectual disability (ID), attention deficit hyperactivity disorder, and others. The underlying causes of most of these pathologies are complex and varied, including prenatal adverse environmental factors and genetic alterations. The development of cutting-edge genomic technology has revealed copy number variations (CNVs) as one of the influential genetic factors contributing to these pathologies.

*CNTN6* is one of the genes in which CNVs have been associated with ID and ASDs^[1–3]^. *CNTN6* encodes neuronal cell adhesion molecules, primarily expressed in the nervous system^[4–6]^. Data obtained from rodent model systems revealed the involvement of *Cntn6* in various processes during brain development, including oligodendrocyte differentiation^[7]^, dendritogenesis^[8]^, axonogenesis^[9]^, and synapse formation^[10,11]^. A comparative analysis of brain cytoarchitecture between wild-type and *Cntn6-*mutant mice disclosed alterations in the numbers of specific neurons in the cortex^[12]^, while the sizes of various brain parts in mutant mice remained unchanged^10^. Loss of Cntn6 also led to behavioral changes in Cntn6-knockout mice, including impaired motor locomotion^[13]^, allocentric navigation^[14]^, and reproductive behavior^[15]^. However, despite the similarity in the early stages of mammalian brain development, the disease manifestations in humans caused by *CNTN6* mutations are more complex, suggesting the possible presence of species-specific functions of this gene. Additionally, the incomplete penetrance of *CNTN6* CNVs^[1,16,17]^ indicates uncertain genotype-phenotype correlations and dramatically complicates the understanding of disease pathogenesis. Consequently, the role of *CNTN6* in human brain development remains unclear.

In this study, we generated cerebral organoids (COs) from patient-specific and genome-edited induced pluripotent stem cell (iPSC) lines with different *CNTN6* genetic variants. We found that the *CNTN6* locus regulates morphogenesis, including lumenization and cell fate determination of radial glial cells (RGCs), during the early stages of human cortical development. Additionally, we demonstrated that *CNTN6* is involved in RGC population maintenance by positively regulating symmetric proliferative division. Bulk transcriptome analysis revealed that loss of *CNTN6* results in gene expression alterations that drive both early and late cortical development and global changes in the signaling pathway landscape. Furthermore, we found that CNTN6 acts through the Notch signaling pathway during early cortical development, interacting with NOTCH1 and NOTCH2 receptors. Finally, CNTN6 may play a role in the nucleocytoplasmic translocation of PAX6 protein through the interaction of the intracellular domain of NOTCH1 with PAX6. Notably, we established that various genetic variants contributed differently to the mutant phenotypes, suggesting that *CNTN6* copy number variation or other structural effects, rather than protein loss alone, may critically modulate early neurodevelopmental outcomes. Thus, our data indicate previously unknown functions of the *CNTN6* locus in the early stages of human cortical development.

## Results

### CNTN6 is expressed in the RGCs during the early stages of human cerebral cortex development

Previous reports demonstrated a peculiar expression pattern of the *Cntn6* gene during rodent cerebral cortex development. It was found that *Cntn6* was expressed at an extremely low level during embryonic development, and its expression gradually increased in the postnatal period, reaching the maximum at the P7 stage^[6]^. During the embryonic period of cerebral cortex development, *Cntn6* expression was first detected in the intermediate zone at the E15.5 stage, then in deep-layer pyramidal cortical neurons at the E17.5 stage^[8]^. Postnatally, *Cntn6* expression was predominantly limited to layer V of the visual cortex^[8,12]^. However, little is known about *CNTN6* expression throughout human cortical development, especially at the embryonic stages. To clarify this issue, we generated COs from a healthy control iPSC line **(Figure S1A)** and analyzed CNTN6 expression immunohistochemically. Initially, we focused on the 45-day stage since COs at this stage contain deep-layer neurons in which CNTN6 expression could be detected according to the rodent model. In COs at 45 days of differentiation, CNTN6 protein was expressed throughout the cortical wall **(Figure 1A).** The highest expression was observed in CTIP2+ deep-layer neurons located in the cortical plate and, surprisingly, in SOX2+ RGCs at the apical surface of the ventricular zone (VZ)-like structures. A similar pattern was obtained in the human embryonic brain at the 8 gestational weeks (GW) **(Figure 1B)**. This unexpected expression in RGCs led to the hypothesis that CNTN6 might also be expressed at earlier stages of development. We found that in the 20-day COs, the signal enrichment was also detected in the apical surface of the VZ-like structures containing predominantly PAX6+ RGCs **(Figure 1B)**. Next, to investigate the temporal dynamics of CNTN6 expression, we performed a western blot analysis of COs at different stages: 20, 30, 45, and 90 days **(Figure 1C)**. The protein level was lowest at day 20, then gradually increased, reaching a maximum at 45 days **(Figure 1D)**. Interestingly, we noted a decreased level in 90 days. It has been previously shown in rats that Cntn6 is expressed in neurons but not in astrocytes and oligodendrocytes^[7]^. Given that gliogenesis during CO development progresses actively after two months and beyond^[18]^, we suggested that the increased percentage of CNTN6-negative non-neuronal cells in later-stage COs may lead to a decreased level of CNTN6 protein.

**Figure 1.**
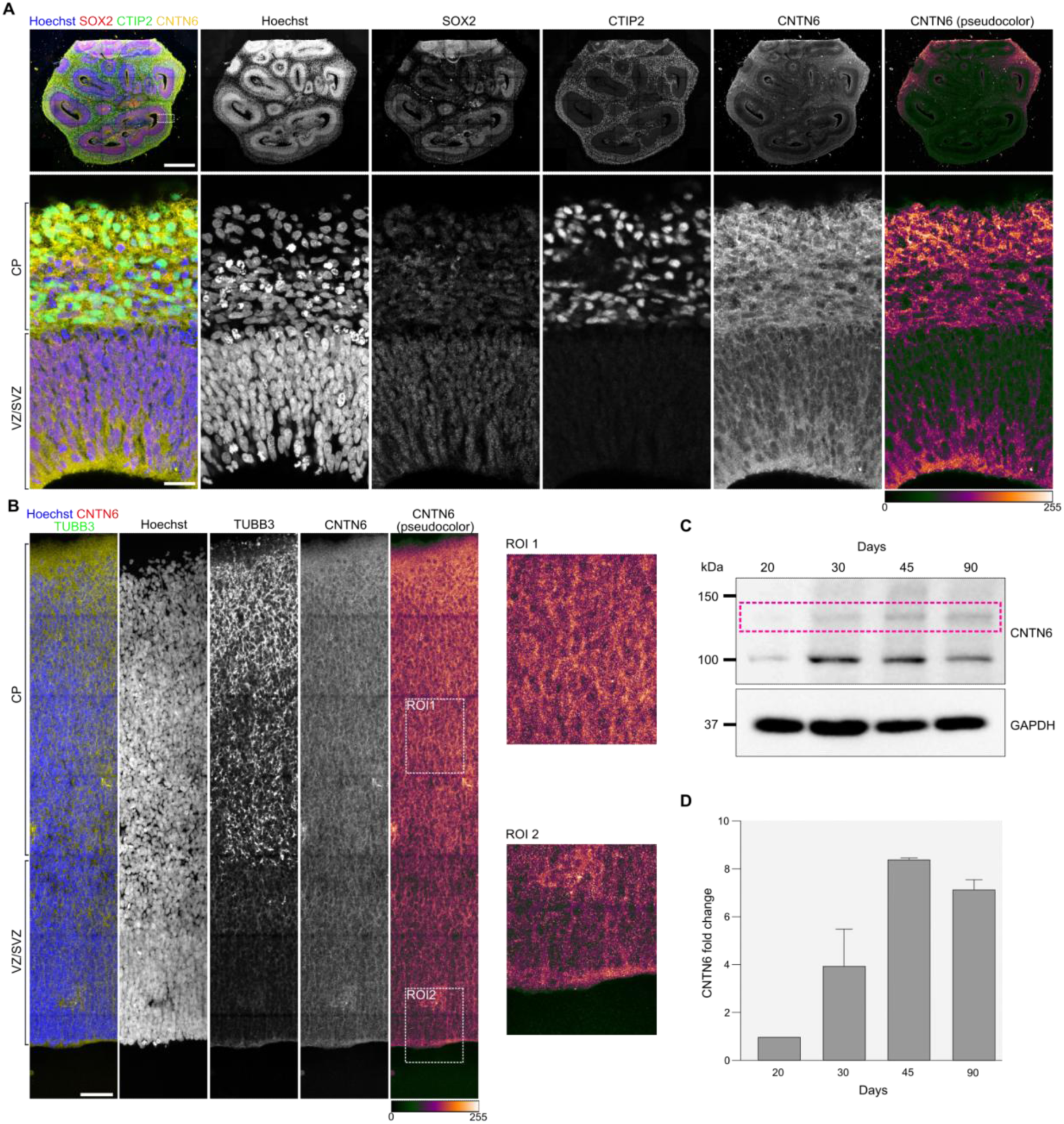
Spatiotemporal expression of CNTN6 during the early stages of human cerebral cortex development. **A, B** Immunohistochemical analysis of CNTN6 expression on day 45 CO (**A**) and human fetal brain at stage GW8 (**B**). Separated channels are presented in grayscale. In addition, a pseudocolor representation of the signal distribution is presented for the CNTN6 channel. The white dotted rectangle labels ROI. CTIP2 and SOX2 mark deep-layer neurons and radial glial cells, respectively (**A**). TUBB3 marks neurons (**B**). VZ – ventricular zone; SVZ – subventricular zone; CP – cortical plate; ROI – region of interest. Scale bar: 500 μm (upper panel **A** and **B**) and 50 μm (bottom panel **A**). **C** Representative western blot results of CNTN6 protein expression in COs at different stages of differentiation. Pink dotted rectangle labels bands corresponding to CNTN6. **D** Quantification analysis of CNTN6 band intensity from western blot normalized to GAPDH. Data is presented as a mean of fold change (n=2 for each stage) ± SD relative to the corresponding day 20 COs.

Additionally, we investigated the СNTN6 expression in two distinct populations of human neural cells: neural stem cells (NSCs), which have similar features to the RGCs, and post-mitotic neurons **(Figure S1C-E)**. Both cell populations were derived from a healthy donor’s iPSC line by dual SMAD inhibition and subsequent terminal differentiation. As expected, the neurons expressed CNTN6, as confirmed by immunohistochemistry **(Figure S1F)** and western blot analysis **(Figure S1G),** thereby corroborating previously published data^[7]^. Surprisingly, we also discovered that NSCs expressed CNTN6, aligning with our CO and fetal brain results.

Collectively, these findings suggest an earlier onset of CNTN6 expression during human cortex development compared to rodents.

### CNVs and knockouts in the *CNTN6* locus lead to disruption of lumenization and RGC cell fate determination in COs

The overwhelming majority of patients with *CNTN6*-related neurodevelopmental disorders have CNVs^[1–3]^, rather than point mutations disrupting gene function. We implied that the type of the genetic variant may be crucial for the severity of the pathology manifestation. To verify this assumption, we assembled a panel of iPSC lines with deletions and knockouts in *CNTN6* **(Figure 2A)**. Several lines from two healthy donors (Control #1 and Control #2) and two sibling patients with a heterozygous ∼400-kb deletion of *CNTN6* (*CNTN6^Δ/+^*#1 and *CNTN6^Δ/+^#*2) were generated and characterized previously^[19–22]^. Subsequently, novel knockout iPSC lines with indel mutations, designated as *CNTN6^Δ/mut^* and *CNTN6^mut/mut^*, were generated by the CRISPR/Cas9 system from patient-derived *CNTN6^Δ/+^*#1 and Control #1 donor-derived iPSC lines, respectively. Detailed information about iPSCs generation, genetic modifications, and characterizations is provided in **Supplementary Table 1 and Figure S2A-E**.

**Figure 2.**
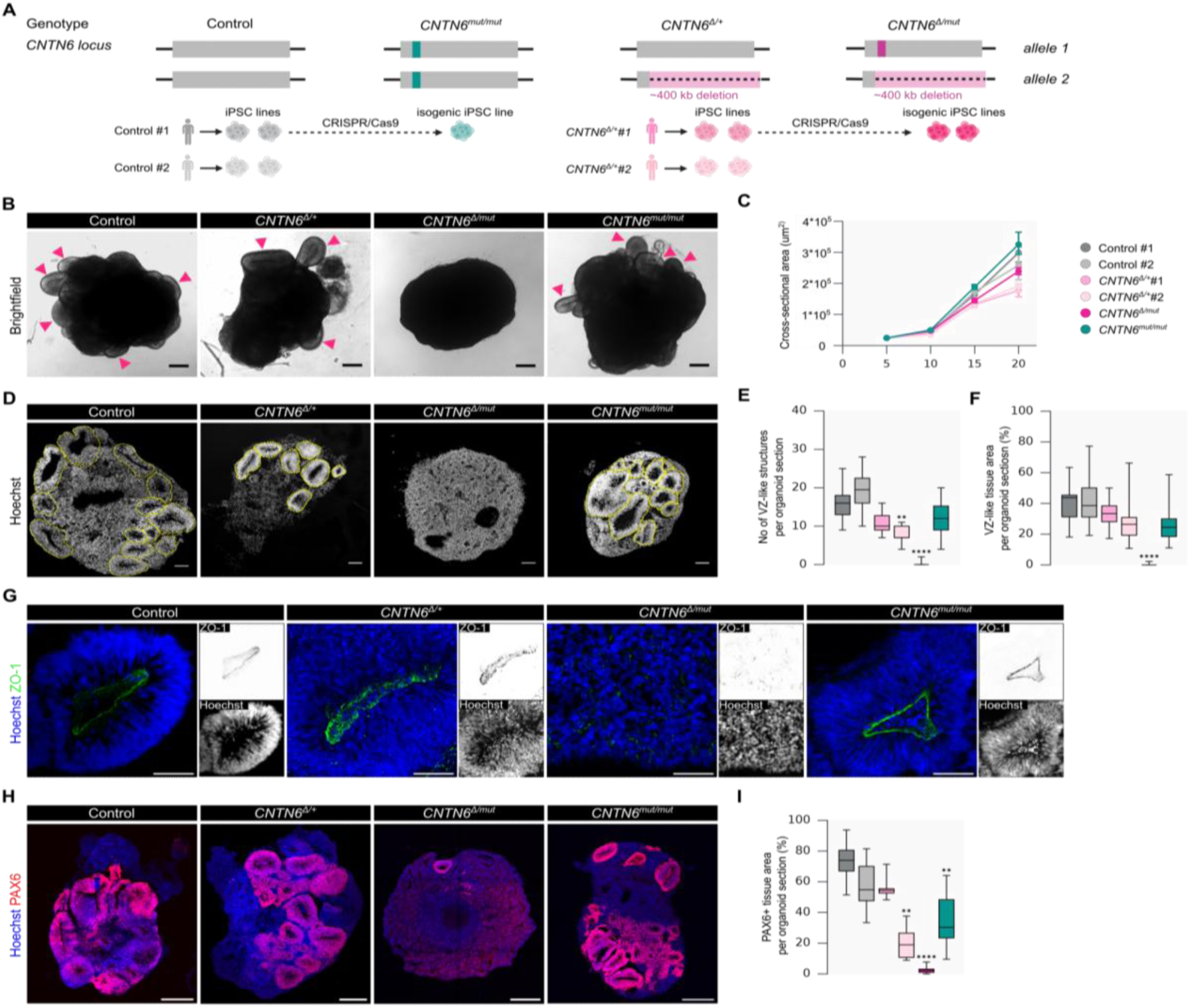
CNV and knockouts in the *CNTN6* gene lead to disruption of VZ-like structure formation during CO differentiation. **A** Schematic depicting the design of different genotypes of iPSCs used for CO generation. **B** Representative brightfield images of COs at day 20. Pink arrowheads label VZ-like structures. Scale bar: 100 μm. **c** Quantification of the cross-sectional area of COs by day. Data points represent the median with 95% confidence intervals (n>40 for each group on any day). **D** Representative fluorescent images of the entire organization of COs at day 20. A yellow dotted line labels VZ-like structures. Scale bar: 100 μm. **E, F** Quantification analysis of morphological parameters of COs: **(E)** the number of VZ-like structures per organoid section and **(F)** the VZ-like tissue area per organoid section. Box plots represent median and quartiles with minimum and maximum values (n>14 for each group; **P = 0.014, ****P < 0.0001; Kruskal–Wallis test with post-hoc Dunn’s test). **G** Representative fluorescent images of ZO-1+ apical membranes. Separated channels are presented in grayscale (Hoechst) and inverted grayscale (ZO-1). Scale bar: 50 μm. **H** Immunohistochemical analysis of PAX6 expression in day 20 COs. Scale bar: 200 μm. **I** Quantification analysis of PAX6+ area distribution in CO sections. Box plots represent median and quartiles with minimum and maximum values (n>5 for each group; **P = 0.0043 (Control #1 versus *CNTN6^Δ/+^#*2), **P = 0.0052 (Control #1 versus *CNTN6^mut/mut^*) ****P < 0.0001; Kruskal–Wallis test with post-hoc Dunn’s test). The color legend for all plots is identical to that in **C**.

To investigate the role of the *CNTN6* locus during the early stage of human cortical development, we generated COs from iPSC lines with different *CNTN6* genotypes and investigated its morphological features. Firstly, we evaluated the growth dynamics of COs during the first 20 days of differentiation **(Figure 2B and Figure S3A,B)**. All iPSC lines formed embryoid bodies (EBs) of comparable size and morphology on day 5. On day 10, also no differences in size were found, except for *CNTN6^Δ/+^*#2, whose EB sizes were reduced by ∼20% compared to Control #1. However, on days 15 and 20, all COs with the 400-kb deletion (*CNTN6^Δ/+^*#1, *CNTN6^Δ/+^*#2, and *CNTN6^Δ/mut^*) exhibited a decrease in size compared to the Control #1 group (**Figure 2C)**. Additionally, we observed alterations in the external morphology of COs at 20 days. *CNTN6^Δ/mut^* COs were relatively spherical and smooth, in contrast to other groups of COs with protrusions containing neuroepithelial buds **(Figure 2B).** At the same time, quantitative analysis revealed reduced surface complexity for *CNTN6Δ/+#1* COs as well **(Figure S3C,D)**. Interestingly, no changes in *CNTN6^mut/mut^* COs were observed in either size or morphology, even after 20 days.

The simplified external morphology of *CNTN6^Δ/mut^*COs suggests potential alterations in their internal organization. To investigate this, we studied lumenization – the formation of VZ-like structures within COs – by assessing its numbers and sizes **(Figure 2D)**. At day 20 of differentiation, the number of VZ-like structures and its occupied area in *CNTN6^Δ/mut^* COs decreased sharply compared to Control #1 **(Figure 2E,F)**.

One of the key conditions for the correct formation of the lumen is the apical-basal polarization of neuroepithelial cells, which occurs at the stage of closure of the neural tube. Polarization is accompanied by the emergence of tight junctions, which are essential for the formation of the apical membrane. We decided to test whether apical membrane formation was impaired in *CNTN6^Δ/mut^* COs. We studied the distribution of polarity junction protein ZO-1 in COs to test it. On day 20, in almost all groups of COs, we detected a lot of VZ-like structures, with a strong specific ZO-1 signal in the apical surface **(Figure 2G)**. In contrast, *CNTN6^Δ/mut^*COs demonstrated a weak ZO-1 signal and an absence of clearly defined apical membranes due to the disorganized distribution of the protein. These findings suggest that compound *CNTN6* variants, including deletion and loss-of-function, impair lumenization during the early stages of human cortical development by disrupting neuroepithelial polarity.

It is known that polarity and cell fate are intricately linked^[23]^. Our data indicate that in *CNTN6^Δ/mut^* COs, neuroepithelial cells lose their ability to polarize. We decided to investigate whether the correct cell fate determination occurs in neuroepithelial cells during the differentiation of COs with different *CNTN6* variants. To determine this, we analyzed the representation and spatial distribution of PAX6+ RGCs in COs **(Figure 2H)**. We detected the presence of PAX6+ cells in COs with different genotypes except *CNTN6^Δ/mut^*. Most of the cells were located within the VZ-like structures. Distinct differences were noted in the qualitative distribution of PAX6+ areas. Quantitative analysis revealed an almost total absence of PAХ6+ cells in *CNTN6^Δ/mut^* COs. For instance, only ∼2% of the organoid section area was PAX6-positive in *CNTN6^Δ/mut^*COs, whereas, in controls, on average, ∼70% of the area was PAX6-positive **(Figure 2I)**. Moreover, decreased levels were also observed for *CNTN6^Δ/+^*and *CNTN6^mut/mut^* COs, with 4 and 2.4-fold change, respectively.

Taken together, the obtained data suggest that genetic *CNTN6* variants disrupt early cortical development through distinct mechanisms: while morphological defects such as impaired lumenization appear to be predominantly driven by CNVs, alterations in RGC identity and fate specification are more closely associated with the loss of CNTN6 protein. Although these two developmental processes are inherently linked—since proper morphogenesis provides the structural context for fate specification and vice versa—our data indicate that different classes of *CNTN6* variants influence early neurodevelopment through somewhat independent molecular pathways. Thus, the phenotypic severity and nature of developmental disruption may depend on the type and molecular consequences of the *CNTN6* variants, highlighting the complexity of *CNTN6*-associated neurodevelopmental disorders.

### CNVs and knockouts in *CNTN6* promote the transition of RGCs to asymmetric neurogenic division

The smaller size of *CNTN6^Δ/mut^* COs and impaired parameters of VZ-like structures led us to guess that the *CNTN6* locus may be involved in the regulation of RGC proliferation. To investigate this, we evaluated the mitotic index and the distribution of symmetrically and asymmetrically dividing cells. Firstly, we examined the proportion of mitotic cells (pH3+) in VZ-like structures **(Figure 3A,B)**. In three groups of COs with the 400-kb deletion, the proportion of pH3+ cells was decreased by 1.9 times for *CNTN6^Δ/+^* #1 and *CNTN6^Δ/mut^*, 2.5 times for *CNTN6^Δ/+^* #2 compared to Control #1 **(Figure 3B)**. These results suggest that the CNVs at the *CNTN6* locus reduce the mitotic output of RGCs, potentially limiting the progenitor pool available for cortical development.

**Figure 3.**
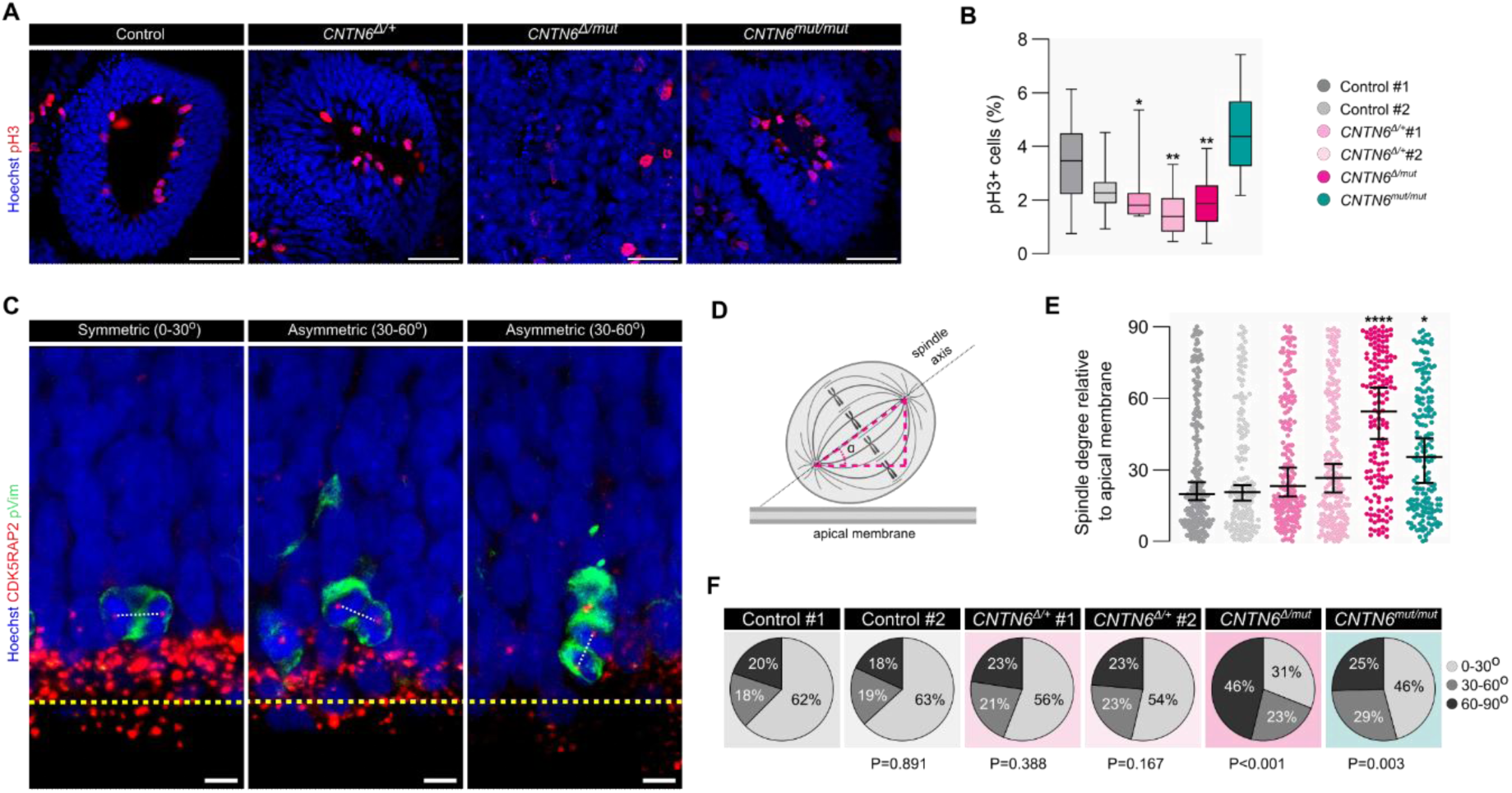
The violation of RGCs proliferation in COs with deletion and knockouts in *CNTN6*. **A** Representative fluorescent images of immunohistochemical analysis of the pH3 expression in COs on day 20. Scale bar: 50 μm. **B** Quantification of the percentage of pH3+cells in COs on day 20. Box plots represent median and quartiles with minimum and maximum values (n>9 organoid sections for each group; *P = 0.0336, **P = 0.0011 (Control #1 versus *CNTN6^Δ/+^#*2), **P = 0.0050 (Control #1 versus *CNTN6^Δ/mut^*); Kruskal–Wallis test with post-hoc Dunn’s test). **C** Representative fluorescent images with examples of the mitotic spindle angle. Yellow and white dotted lines label the apical surface and the spindle, respectively. Scale bar: 5 μm. **D** Schematic of spindle angle measurements. **E, F** Quantification of the mitotic spindle angle. Scatter-dot plot **(E)** represents median and individual values with confidence interval (n>158 cells for each group; *P = 0.0119, ****P < 0.0001; Kruskal–Wallis test with post-hoc Dunn’s test). **F** Circle chart represents the distribution of symmetric (spindle angle 0-30°) and asymmetric (spindle angle 30-60° and 60-90°) divisions (n>158 cells for each group; P-values are indicated on figure; Chi-square test).

Next, we assessed spindle orientation in dividing RGCs, which may reflect the distribution of symmetric proliferative and asymmetric neurogenic divisions. Notably, an imbalance in the distribution of these types of divisions can lead to premature neurogenesis^[24–26]^. We analyzed the mitotic spindle angle to determine whether a similar defect is present in COs with *CNTN6* mutations **(Figure 3C-E).** The mitotic spindle orientation was predominantly horizontal in control COs with a median angle of ∼20° **(Figure 3D)**. In contrast, dividing cells in knockout *CNTN6^Δ/mut^*and *CNTN6^mut/mut^* COs demonstrated more frequent oblique or vertical spindle orientation and significantly increased median angles of ∼50° and 40°, respectively **(Figure 3E)**. Quantitative analysis of distribution showed that in the case of mutant *CNTN6^Δ/mut^* and *CNTN6^mut/mut^* COs, the proportion of asymmetrically dividing cells increased by 32% and 17% respectively, compared to the Control group **(Figure 3F).** Although spindle angle measurements may not fully and precisely distinguish between division types, they can offer valuable insights into changes in division patterns and potential shifts in cell fate specification. Interestingly, the reduction in mitotic activity was observed only in COs with the 400-kb deletion, not in those with loss of CNTN6 protein alone. This suggests that CNVs may uniquely affect RGC proliferation, likely through additional regulatory mechanisms beyond protein loss. In contrast, primarily the loss of the CNTN6 protein influenced the balance between symmetric and asymmetric divisions. These findings indicate that various types of *CNTN6* genetic variants may have different molecular effects, influencing human cortical development through distinct mechanisms.

### The phenotype of COs with *CNTN6* deletion is partially rescued by the overexpression of the CNTN6 protein and the chimeric organoid approach

To understand the role of CNTN6 protein in lumenization and cell fate determination, we performed rescue phenotype experiments. Firstly, we generated a genetically modified *CNTN6^Δ/mut_AAVS1-CNTN6-GFP^* iPSC line with doxycycline-inducible overexpression of CNTN6 and GFP **(Figure 4A, Figure S4A,B)**. The integration of the genetic construct into the *AAVS1* locus and its functionality were validated by PCR **(Figure S4C)**. Next, we generated COs with time-controllable CNTN6 overexpression from day 15 of differentiation **(Figure 4B-D)** and analyzed its morphology. We found that the internal organization of COs was partially restored as the number of VZ-like structures increased significantly **(Figure 4E,F).** However, the sizes of mutant COs were still significantly smaller than the Control #1 COs **(Figure S4D,E)**. We also noted that ZO-1 expression was partially restored in the apical surface of the VZ-like structures **(Figure 4G)**, but cells inside the structures did not express PAX6 protein **(Figure 4H)**. Thus, our findings indicate that overexpression of *CNTN6* leads to a partial rescue of the *CNTN6^Δ/mut_AAVS1-CNTN6-GFP^* mutant phenotype in lumenization, but not in cell fate determination. Given that CNTN6 is typically expressed at very low levels, we hypothesized that partial rescue of the mutant phenotype could be a side effect of protein overexpression. Moreover, by overexpression, we largely overstimulated it as a cell-autonomous factor. However, several studies have shown that CNTN6 acts as a ligand in both cis-^[27]^ and trans-interactions^[7]^ with receptors and is thus also an extracellular, i.e., non-cell-autonomous factor.

**Figure 4.**
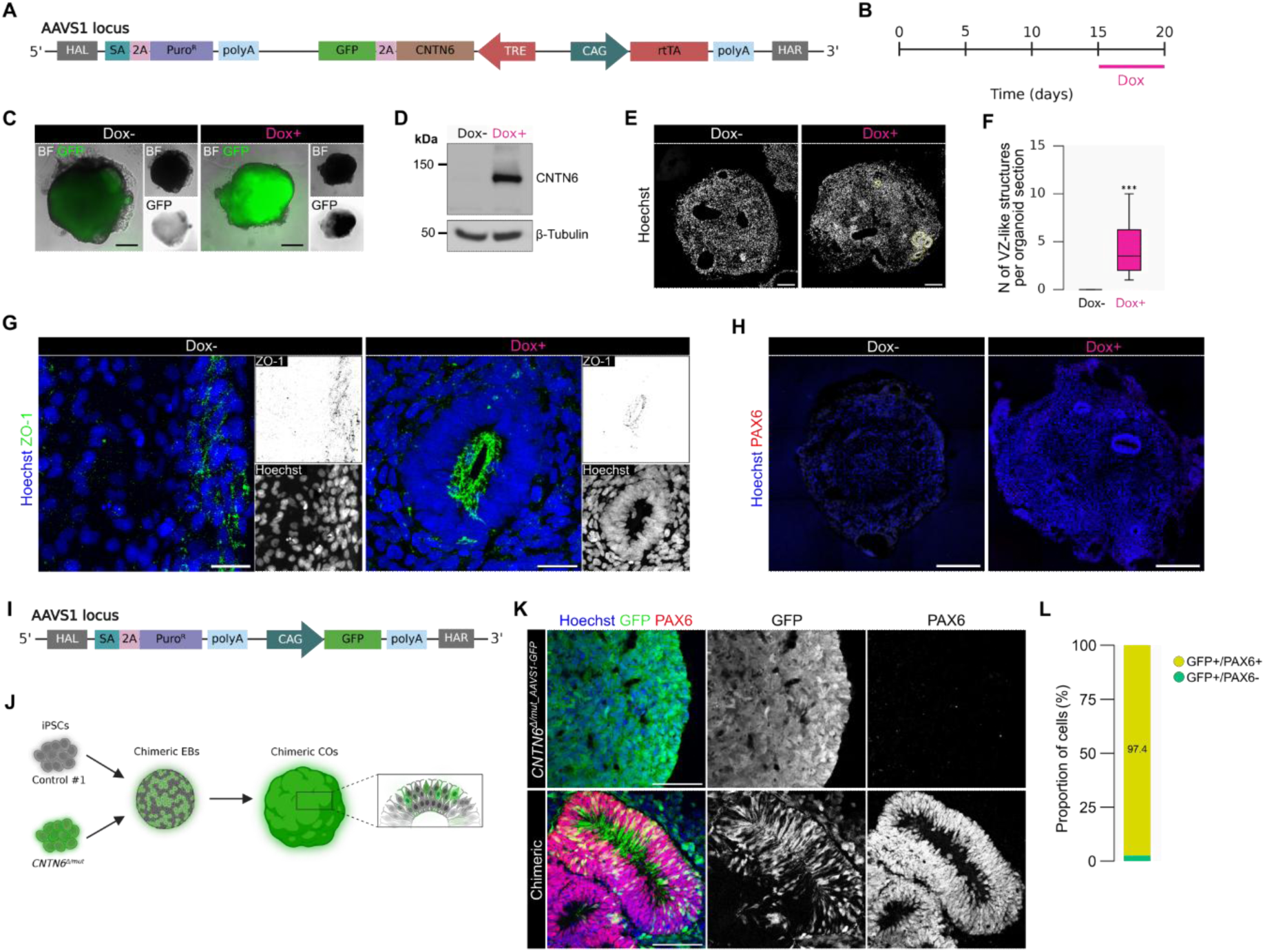
Partial rescue of phenotype in *CNTN6^Δ/mut^* COs by *CNTN6* overexpression and chimeric organoid approach. **A** Simplified schematic representing the modification of the *AAVS1* locus in iPSC line *CNTN6^Δ/mut^*for doxycycline-inducible overexpression of CNTN6 and GFP. HAL/HAR - homology arms left/right; SA - splice acceptor; TRE - Tet response element; rtTA - reverse tetracycline-controlled transactivator. **B** Schematic depicting design of experiment conditions used for CO differentiation with doxycycline-inducible expression of CNTN6 and GFP. **C** Representative combined brightfield and fluorescent images of doxycycline-treated (Dox+) and control (Dox-) COs at day 20. A separated channel with GFP is presented in an inverted grayscale. Scale bar: 200 μm. **D** Representative western blot results of doxycycline-inducible CNTN6 protein expression in COs at day 20. **E** Representative fluorescent images of the entire organization of COs at day 20 with doxycycline-inducible CNTN6 overexpression. A yellow dotted line labels VZ-like structures. Scale bar: 200 μm. **F** Quantification analysis of the number of VZ-like structures per CO section. Box plots represent median and quartiles with minimum and maximum values (Dox-n=8, Dox+ n=12; ***P < 0.000148; Mann-Whitney U test). **G** Representative fluorescent images of ZO-1+ apical membranes. Separated channels are presented in grayscale (Hoechst) and inverted grayscale (ZO-1). Scale bar: 25 μm. **H** Immunohistochemical analysis of PAX6 expression in COs at day 20 with doxycycline-inducible CNTN6 overexpression. Scale bar: 250 μm. **I** Simplified schematic representing the modification of the *AAVS1* locus in iPSC line *CNTN6^Δ/mut_AAVS1_GFP^ f*or constitutive overexpression of GFP. **J** Simplified schematic depicting generation of chimeric COs from iPSCs with *CNTN6^Δ/mut_AAVS1_GFP^*and Control #1 genotypes. **K** Immunohistochemical analysis of PAX6 expression in chimeric COs and *CNTN6^Δ/mut_AAVS1_GFP^* at day 20. Separated channels are presented in grayscale. Scale bar: 100 μm. **L** Quantification analysis of the PAX6 expression in chimeric and COs. GFP+ cells located within VZ-like structures were included in the analysis. Scattered bar plots represent the percentage of GFP+/PAX6+ and GFP+/PAX6-cells in chimeric COs (n=991cells).

To overcome these potential limitations, we modified the experimental design to ensure that *CNTN6* is expressed at a physiological level and that it can be implemented as a non-cell-autonomous factor. We hypothesized that mutant cells surrounded by normal cells expressing CNTN6 would be rescued and contribute to VZ-like structures. To address this issue, we generated chimeric COs by mixing cells with different genotypes. We created a genetically modified *CNTN6^Δ/mut_AAVS1_GFP^* iPSC line expressing GFP (**Figure 4E and Figure S4A,B,F**) to label mutant cells inside COs. Next, we utilized a mixture of Control #1 and GFP-labeled *CNTN6^Δ/mut_AAVS1_GFP^* cells to generate chimeric COs (**Figure 4I**). Notably, we observed that chimeric COs formed well-organized VZ-like structures with a clear contribution of GFP+ *CNTN6^Δ/mut_AAVS1_GFP^* cells (**Figure 4J, Figure S4G**). Moreover, 97% of these mutant cells, located within VZ-like structures, exhibited restored PAX6 expression **(Figure 4L)**. This observation supports a non-cell-autonomous mechanism, whereby the presence of wild-type cells in the chimeric environment facilitates the rescue of PAX6 expression in neighboring mutant cells.

Together, these results indicate that the CNTN6 protein plays a crucial role in the lumenization and cell fate determination during the early stages of human cortical development, operating through non-cell autonomous mechanisms.

### CNVs and knockouts in *CNTN6* lead to alterations of gene expression and global changes in the signaling pathway landscape in COs

Previous studies have shown that *CNTN6* may act through different signaling pathways depending on the cell type and process^[7,27–30]^. However, nothing is known about its molecular mechanism of action in RGCs during the early stages of human brain development. To address this issue, we performed bulk RNA-seq on 45-day COs (**Figure 5A**) since, at this stage, both populations of *CNTN6*-expressing cells, post-mitotic neurons, and RGCs are present. A comparison of *CNTN6^mut/mut^*and Control #1 COs revealed 1020 differentially expressed genes (DEGs) **(Supplementary Table 2)**, while 860 DEGs were identified for *CNTN6^Δ/mut^* COs **(Supplementary Table 3)**. Gene ontology (GO) analysis for each DEG set revealed several categories related to brain development **(Figure S5A,B, Supplementary Table 4,5)**. Next, we determined the general effect of different *CNTN6* mutations by crossing both sets of DEGs and identified 175 genes whose expression was altered in both mutant types of COs **(Supplementary Table 6)**. Interestingly, this set of DEGs was enriched for the GO terms, which display processes previously known to involve *CNTN6* gene, for instance: “chemical synaptic transmission” (GO:0007268), “axon development” (GO:0061564), “regulation of oligodendrocyte differentiation” (GO:0048713), and “dendrite” (GO:0030425) (**Figure 5B, Supplementary Table 7)**. Moreover, among the DEGs, we observed several categories that were consistent with our experimental results and reflected morphological changes in *CNTN6* mutant COs, including “cell fate commitment” (GO:0045165), “neural tube formation” (GO:0021915), and “neural precursor cell proliferation” (GO:2000178) (**Figure 5C**).

**Figure 5.**
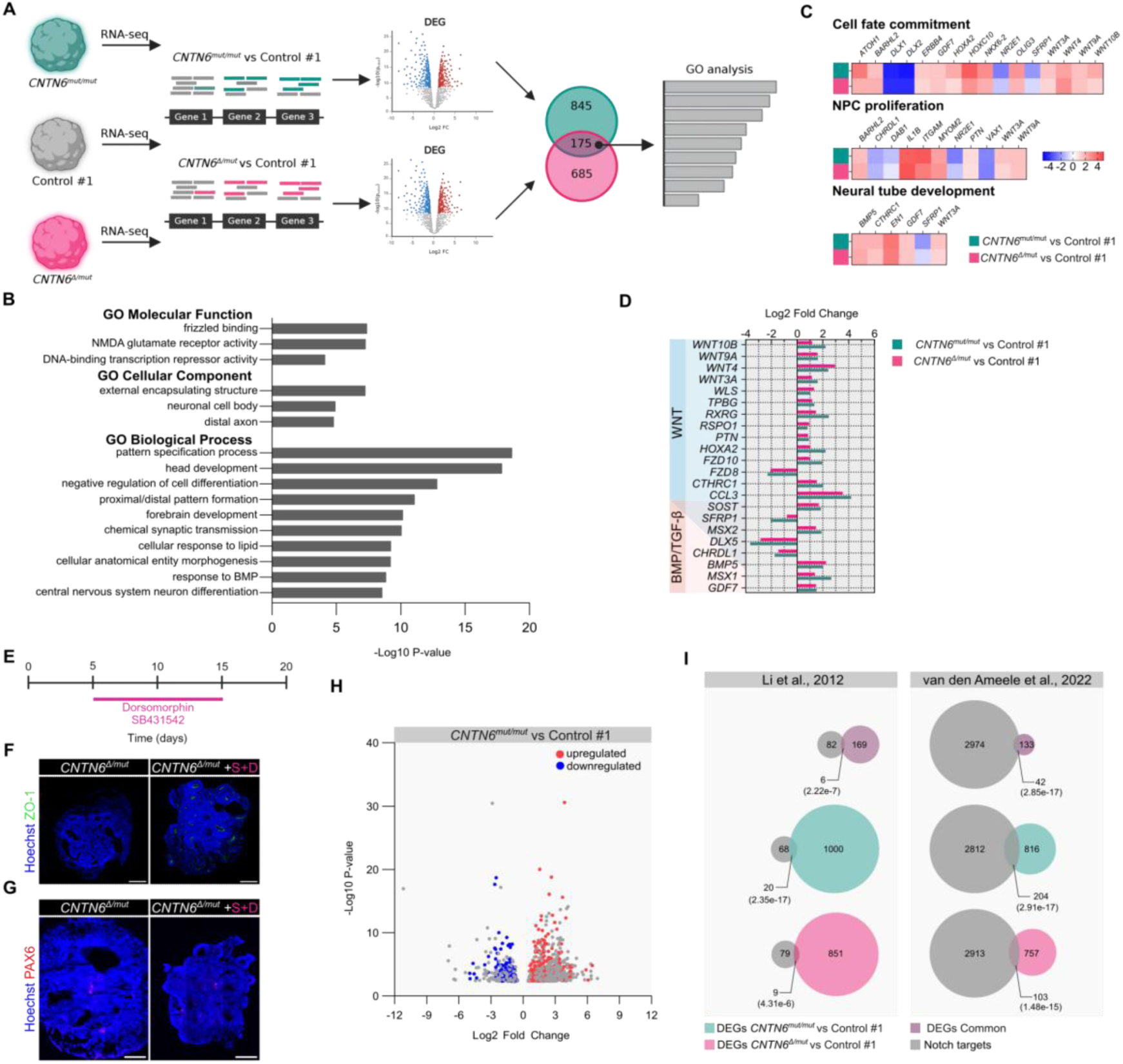
Comparative gene expression analysis of COs with deletion and knockouts in the *CNTN6* locus. **A** Schematic depicting the design of bulk RNA-seq analysis. **B** GO overrepresentation tests using Metascape (https://metascape.org/) for common DEGs identified from CO RNA-seq are shown. The bar chart represents -Log10 P-values for top-ranked GO terms. Metascape uses the hypergeometric test for enrichment P-value calculation. **C** Heatmaps for DEGs from GOs associated with the *CNTN6* mutant phenotype of COs. **D** Expression of DEGs associated with WNT and BMP/TGFβ signaling pathways identified by GO-analysis on the common DEG set. The bar chart represents Log2 Fold Change. Pink and green colors represent two DEG sets, “*CNTN6^Δ/mut^* vs Control #1” and “*CNTN6^mut/mut^* vs Control #1”, respectively. **E** Schematic depicting the design of the experiment conditions used for COs differentiation with inhibition of BMP/TGFβ signaling pathway. **F,G** Immunohistochemical analysis of ZO-1 and PAX6 expression in day 20 COs under inhibition of BMP/TGFβ signaling pathway. **H** Volcano plot represents gene expression changes (Log2 Fold Change) in *CNTN6^mut/mut^* compared to control COs. Colored dots indicate Notch target genes. Blue and red dots label downregulated and upregulated, respectively. **I** Scaled Venn diagrams represent the enrichment of Notch downstream targets in DEG sets. The P-values were calculated using a hypergeometric test.

Further, we focused on the GOs related to signaling pathways. Brain development is known to be controlled by multiple signaling cascades, including the WNT, GSK, BMP/TGF-β, and Notch pathways, among others. Among them, GO analysis revealed enrichment in categories related to WNT and BMP/TGF-β signaling pathways **(Supplementary Table 6, 7)**. Most of the DEGs (*WNT3A, WNT4, WNT9A, WNT10B,* and *BMP5*) from these ontologies were upregulated (**Figure 5D**). This may indicate aberrant activation of these signaling pathways and their contribution to the formation of the *CNTN6* mutant phenotype. We hypothesized that modulation of the activity of these signaling pathways may lead to partial rescue of the CO mutant phenotype. To test this assumption, we selected the BMP/TGF-β signaling pathway. Since most of the genes in this category were upregulated, inhibition with small molecules (Dorsomorphin and SB431542) was chosen as a modulation strategy (**Figure 5E**). Treatment of *CNTN6^Δ/mut^* COs with the inhibitors resulted in the appearance of VZ-like structures (**Figure 5C**) and the formation of an apical membrane (**Figure 5F).** However, the expression of PAX6 was not restored (**Figure 5G**). These data indicate that modulation of the BMP/TGF-β signaling pathway results in a partial rescue of the mutant phenotype, manifested by the resumption of lumenization, but not cell fate determination.

Notably, several Notch pathway-related genes were also identified among the DEGs in the *CNTN6^mut/mut^* group **(Supplementary Table 2)**. For instance, there were transcriptional coactivator *MAML3,* gamma-secretase subunit *PSENEN*, and key downstream target *HES1.* It is known that the Notch pathway plays a crucial role in cortex development and targets a lot of genes^[31,32]^. Moreover, it has been shown in rodents that Cntn6 may act as a non-canonical ligand of the Notch1 receptor and lead to subsequent cascade activation ^[7]^. Thus, we decided to investigate whether the DEGs were enriched for Notch downstream targets. To reveal this, we compared the DEGs with the lists of Notch target genes from previously published papers^[31,32]^. It turned out that all groups of DEGs were significantly enriched with Notch target genes from both lists (P-value <10^-5^) (**Figure 5H, I**). These results indicate large-scale changes in gene expression in both types of *CNTN6* knockout COs, regulated by the Notch signaling pathway.

Overall, our data reveal a global perturbation of gene expression and disruption of the signaling pathway landscape in *CNTN6* mutant COs.

### Inhibition of the Notch signaling pathway partially reproduces the *CNTN6* mutant phenotype

Taking into account that in rodents, Cntn6 functions as a non-canonical ligand for the Notch1 receptor^[7]^ and its loss in the COs results in the large-scale expression changes of Notch signaling downstream targets, we suggested that CNTN6 in RGCs may act through this pathway. Consequently, inhibition of the Notch pathway in normal COs might phenocopy the *CNTN6* mutant COs. To test this hypothesis, we examined the morphological features of Control #1 COs under conditions of impaired Notch signaling using the γ-secretase inhibitor DAPT, which prevents the cleavage of Notch receptors, thereby blocking signal transmission and activation of the pathway. COs were treated with DAPT from day 7 of differentiation **(Figure 6A),** as this stage marks the onset of cell commitment to the neuroectoderm, a process in which Notch pathway activity has been previously shown to play a crucial role^[33–35]^. The reduction of Notch signaling in COs was validated by a lower level of NOTCH1 intracellular domain (NICD1) release and its downstream target, HES1. **(Figure 6B).** Firstly, we estimated the dynamics of CO growth. On days 15 and 20, DAPT-treated COs displayed a decreased size compared with control DMSO-treated COs **(Figure 6C).** The internal morphology of DAPT-treated COs was also altered. For instance, we observed a decreased number of VZ-like structures in DAPT-treated COs by ∼14 times **(Figure 6E,F)**. Moreover, the area occupied by VZ-like structures was reduced by 5.5 times in DAPT-treated COs **(Figure 6G)**, as well as its thickness by ∼1.6 times **(Figure 6G)**. These data demonstrate phenotypic similarities between *CNTN6^Δ/mut^* and DAPT-treated COs, manifested by impaired lumenization.

**Figure 6.**
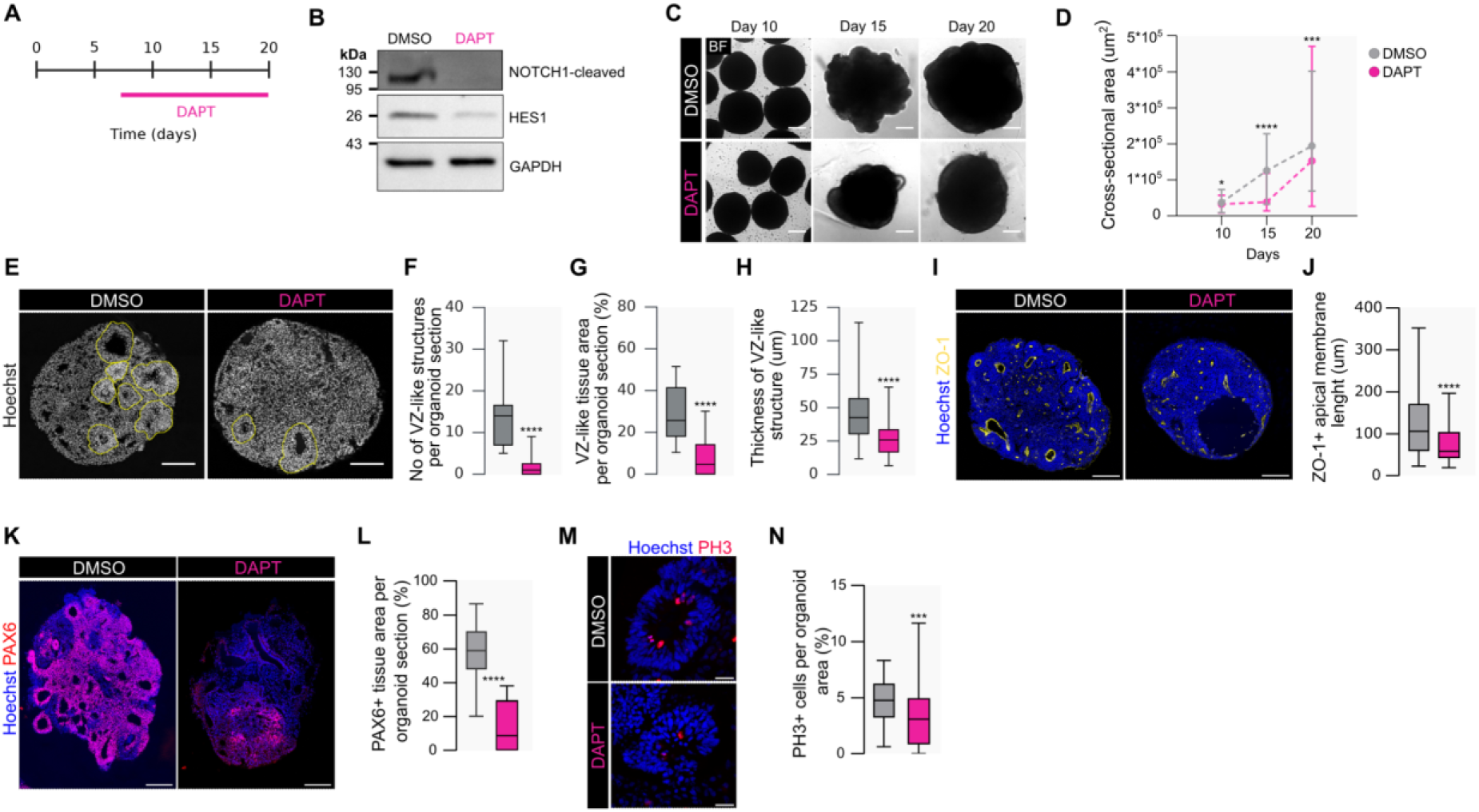
Inhibition of the Notch signaling pathway partially phenocopied *CNTN6^Δ/mut^ COs*. **A** Schematic depicting the design of the experiment conditions used for COs differentiation under DAPT treatment. DMSO was used as a control. **B** Representative western blot results of HES1 protein downregulation and inhibition of NICD1 releasing in COs under DAPT treatment. **C** Representative brightfield images of control and DAPT-treated COs at days 10, 15, and 20. Scale bar: 100 μm. **D** Quantification analysis of the cross-sectional area of DAPT and DMSO-treated COs at different stages of differentiation. Data points represent the median with minimum and maximum values (n>90 for each group on any day; *P=0.0174, ***P=0.0003, ****P<0.0001; Mann-Whitney test). **E** Representative images of the internal organization of DAPT and DMSO-treated COs. VZ-like structures are labeled by the yellow dotted line. Scale bar: 200 μm. **F-H** Quantification analysis of No of VZ-like structures per CO section **(F)**, VZ-like tissue **(G),** and thickness of VZ-like structures **(H)** in DAPT and DMSO-treated COs at 20 days. Box plots represent median and quartiles with minimum and maximum values (n>15 organoid sections for **F** and **G**, n>108 VZ-like structures for **H**; ****P<0.0001; Mann-Whitney test). **I** Immunohistochemical analysis of ZO-1 expression in day 20 COs. Scale bar: 200 μm. **J** Quantification analysis of ZO-1+ length of the apical membrane. Box plots represent median and quartiles with minimum and maximum values (n>252 VZ-like structures for each group; ****P < 0.0001; Mann-Whitney test). **K** Immunohistochemical analysis of PAX6 expression in day 20 COs. Scale bar: 200 μm. **L** Quantification analysis of PAX6+ area distribution in CO sections. Box plots represent median and quartiles with minimum and maximum values (n>12 organoid sections for each group; ****P < 0.0001; Mann-Whitney test). **M** Representative fluorescent images of immunohistochemical analysis of the pH3 expression in COs on day 20. Scale bar: 25 μm. **N** Quantification of the percentage of pH3+cells in COs on day 20. Box plots represent median and quartiles with minimum and maximum values (n>56 VZ-like structures for each group; ***P = 0.001; Mann-Whitney test)

However, DAPT-treated COs retained the ability to form a ZO-1+ apical membrane, unlike *CNTN6^Δ/mut^* COs, though their size was reduced relative to DMSO-treated COs **(Figure 6I,J)**. This finding may indirectly suggest that Notch signaling promotes lumenization, but not as a primary factor. On the contrary, we found that inhibition of Notch signaling led to a dramatic reduction of the CO area occupied by PAX6+ cells by 6.8 times **(Figure 6K,L)**. Notably, selective shRNA-mediated knockdown of *NOTCH1* recapitulated DAPT-treated COs phenotypes **(our unpublished results)**. These data indicate that the disruption of RGC cell fate determination in DAPT-treated COs phenocopies the *CNTN6^Δ/mut^* COs and are consistent with the data supporting a role for Notch signaling in RGC cell fate determination^[36]^. Finally, we examined the proliferation of RGCs as a parameter altered in *CNTN6^Δ/mut^*. The mitotic index in DAPT-treated COs was also affected: the proportion of pH3+ cells decreased by ∼1.5 times **(Figure 6M)**. In summary, our data show that the disruption of Notch signaling in COs results in a phenocopy of most of the aberrations found in *CNTN6* mutant COs, indirectly supporting the significance of CNTN6-dependent Notch pathway activation in RGCs during human cortical development.

### CNTN6 interacts with NOTCH receptors and may contribute to PAX6 cytoplasmic-nuclear translocation via the Notch signaling pathway

A critical step in modulating the Notch signaling pathway is the interaction of Notch receptors with ligands. In rodents, Cntn6 has been demonstrated to interact with the Notch1 receptor, leading to its activation^[7]^. Consequently, we questioned whether human CNTN6 exhibits a similar affinity for NOTCH receptors. We focused on NOTCH1 and NOTCH2 due to their role in RGC maintenance during mammalian cortical development^[36,37]^. To establish this, we examined protein-protein interactions between full-length human CNTN6-3xHA and FLAG-tagged NOTCH1 and NOTCH2 extracellular domains (NECD1 and NECD2, respectively). As a positive control, we used the canonical ligand DLL1-3xHA. Due to the extremely low endogenous expression level of CNTN6 in COs, we used HEK293T for overexpression.

Reciprocal co-immunoprecipitation experiments demonstrated that CNTN6 interacted with both NECD1 and NECD2 **(Figure 7A,B)**. To confirm the activation of Notch signaling following the interaction of CNTN6 with NOTCH receptors, we overexpressed CNTN6 and DLL1 in human NSCs. Semi-quantitative western blot analysis revealed an increased level of downstream target HES5 by 2-3 times mediated by the overexpression of both the canonical ligand DLL1 and CNTN6 **(Figure 7C,D)**. Thus, our results confirm that CNTN6 activates Notch signaling in NSCs.

**Figure 7.**
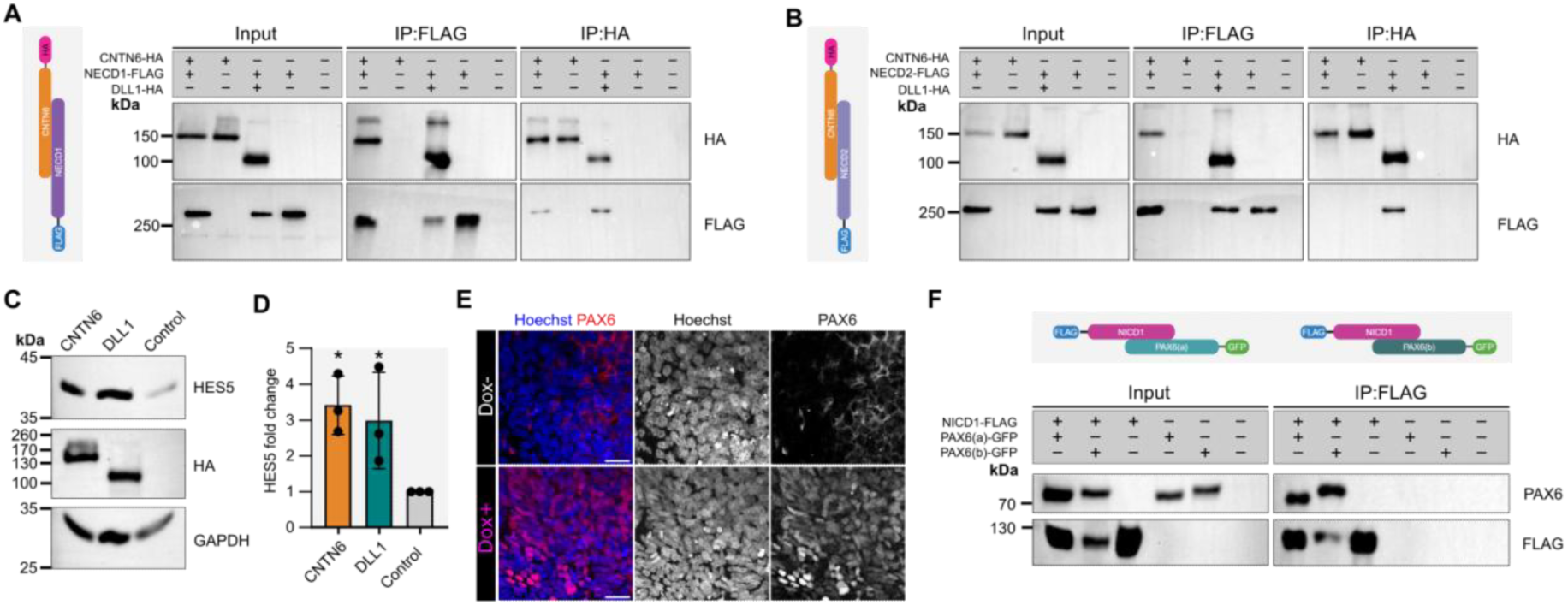
CNTN6 functionally interacts with NOTCH receptors and modulates the subcellular localization of PAX6 protein. **A,B** Co-immunoprecipitation of CNTN6-3xHA with NECD1-FLAG **(A)** and NECD2-FLAG **(B)** confirms the interaction between the proteins in reciprocal reaction. **C** Representative western blot results of HES5 expression in NSCs after 48 h of overexpression of CNTN6 and DLL1. **D** Quantification analysis of HES5 band intensity from western blot normalized to GAPDH. Data is presented as a mean of fold change with SD (n=3; *P= 0.0349 (CNTN6-HA), *P= 0.0368 (DLL1-HA); ordinary one-way ANOVA test with post-hoc Holm-Sidak’s test). **E** Immunohistochemical analysis of *CNTN6^Δ/mut_AAVS1-CNTN6-GFP^* iPSC line at 11 days of differentiation. Separated channels are presented in grayscale. Scale bar: 25 μm. **F** Co-immunoprecipitation of PAX6(a)-GFP and PAX6(b)-GFP with NICD1-FLAG confirms the interaction between proteins.

The Notch signaling pathway regulates numerous downstream targets, including the transcription factor PAX6^[32]^. Immunohistochemical and transcriptomic analyses revealed a significant reduction in PAX6 expression in CNTN6-deficient COs, both *CNTN6^Δ/mut^*and *CNTN6^mut/mut^.* Given this observation, we hypothesized that Notch signaling mediates the molecular link between CNTN6 and PAX6. To directly assess whether CNTN6 influences PAX6 levels via Notch signaling, we hypothesized that CNTN6 overexpression would restore PAX6 expression. However, our data from the phenotype rescue experiment showed that overexpression of *CNTN6* does not lead to PAX6 appearance in COs.

Given the inherent variability of organoid models due to their stochastic self-organization, we sought to validate these findings in a more controlled system. To this end, we employed dual SMAD inhibition, a well-established method that enables the efficient (up to 95%) generation of PAX6⁺ neuroectodermal cells^[38]^. Using the *CNTN6^Δ/mut_AAVS1-CNTN6-GFP^* iPSC line, we examined PAX6 expression at 11 days of differentiation since most cells at this stage are typically PAX6+. In doxycycline-treated *CNTN6*-overexpressing cells, we identified clusters of cells with nuclear PAX6 localization, whereas in control *CNTN6^Δ/mut_AAVS1-CNTN6-GFP^*cells, we observed dispersed regions with its cytoplasmic subcellular localization **(Figure 7E).**

These observations led us to hypothesize that NICD, beyond its well-established role as a transcriptional co-activator, may also function as a molecular shuttle facilitating PAX6 nuclear translocation. To check this hypothesis, we performed co-immunoprecipitation assays. Due to the crucial roles of both PAX6(a) and PAX6(b) isoforms in human neuroectodermal fate specification, we co-expressed each isoform fused to GFP alongside NICD1 (804 aa) tagged with 1xFLAG. Co-IP analysis revealed direct interactions between NICD1 and both PAX6(a) and PAX6(b) **(Figure 7F).**

Together with RNA-seq data from CNTN6-deficient COs, these findings suggest that CNTN6 protein loss disrupts neural progenitor development by downregulating PAX6 expression and putatively impairing its intracellular trafficking, thereby perturbing early fate specification.

## Discussion

Our results suggest that the *CNTN6* locus has a pleiotropic effect on the early development of the human cerebral cortex, influencing tissue morphogenesis, cell identity, and the proliferation of RGCs. We demonstrated that different variants in *CNTN6* disrupt the formation of VZ-like structures and reduce the proportion of PAX6+ RGCs. Additionally, the *CNTN6* locus is involved in RGC proliferation by positively regulating symmetrical cell divisions. The functions of *CNTN6* in early human cortical development are partially mediated through the activation of the Notch signaling pathway. Furthermore, we observed that CNVs in *CNTN6* manifest more robust phenotypic effects compared to the loss-of-function mutations. *CNTN6* demonstrates an early onset of expression during human cortical development, in contrast to its later expression observed during rodent embryogenesis^[6–8]^. Previous studies have shown that cortex *CNTN6* expression was mainly restricted to layer V of the visual cortex^[6,8,12,27]^, where deep-layer pyramidal neurons are located. Consistent with these findings, we observed *CNTN6* expression in deep-layer CTIP2-positive neurons located in the cortical plate of COs. However, we also found expression of *CNTN6* in RGCs. These observations suggest that *CNTN6* can play a broader role in human cortical development, encompassing not only the maturation of postmitotic neurons but also the proliferation, self-renewal, and differentiation of neural progenitors.

According to our data, deletion in the *CNTN6* gene impairs lumenization at the early stages of human cortex development. Cerebral cortical morphogenesis, including neural tube closure, migration, axon, and dendrite guidance, is orchestrated by a set of adhesion molecules that drive complex cell-cell and cell-extracellular matrix interactions^[39,40]^. As a member of the Immunoglobulin superfamily cell adhesion molecules, *CNTN6* also plays an important role in neurite outgrowth^[13,41,42]^, as well as axon^[9,27,43]^ and dendrite navigation^[8]^. However, its involvement in neural tube closure remains unexplored. Our data suggest that *CNTN6* may contribute to establishing apical-basal cell polarity in neural progenitor cells, a process required for proper neural tube closure. The formation of apical-basal polarity during neural development depends on the assembly and regulation of specialized junctional complexes, such as tight and adherens junctions, which include the apically localized protein ZO-1^[44,45]^. Its loss leads to embryonic tissue disorganization and causes lethality^[46]^. We hypothesize that *CNTN6* positively regulates *ZO-1* expression during early human cortex development. Disruption of ZO-1 production and its altered spatial distribution in *CNTN6*-deficient COs, along with its recovery upon *CNTN6* overexpression, support this hypothesis. Interestingly, *CNTN6*-dependent regulation of *ZO-1* expression has also been reported in endothelial cells^[47]^, although the molecular mechanism underlying this regulation remains unclear and warrants further investigation.

In addition to morphogenesis, tissue patterning is essential for shaping the human cerebral cortex, in which the correct determination of cell fate is one of the fundamental aspects of this process. Our findings indicate that variants in *CNTN6* impair RGC fate specification, as reflected in reduced *PAX6* expression. Given that *PAX6* is a master regulator governing neural patterning^[48–50]^, including cell fate determination^[51,52]^, its dysregulation has profound developmental consequences. One of the notable outcomes in *CNTN6*-deficient COs is a shift in RGC division from symmetric proliferative to asymmetric neurogenic modes. This change may alter the balance of progenitor cell populations and disrupt proper cortical development, consistent with the previous findings in Pax6-deficient mouse models^[53,54]^. But, in contrast, while *Pax6* deficiency in the developing mouse cortex is also associated with increased progenitor proliferation^[55–57]^, we observed a decreased proliferative capacity of RGCs in *CNTN6* mutant COs. This effect was present only in COs harboring 400-kb deletion in the *CNTN6* gene, suggesting a distinct molecular mechanism beyond simple CNTN6 protein deficiency. Unraveling this mechanism may provide key insights into the etiology of microcephaly in some patients with both deletions and duplications of the *CNTN6* gene ^[1,17,58,59]^, potentially leading to better diagnostic and therapeutic approaches for these patients. The simultaneous disruption of morphogenesis and tissue patterning in *CNTN6*-mutant COs raises an important question: is one of these defects a direct consequence of the other, or are they independent? Interestingly, both processes are governed by the same core signaling pathways^[36]^ that are intricately cross-regulated^[60]^, potentially linking these defects through shared molecular mechanisms. Transcriptomic analysis of CNTN6-deficient СOs revealed broad changes in WNT, BMP/TGF-β, and NOTCH signaling, highlighting their potential involvement in mediating CNTN6-dependent cortical development. However, although CNTN6 acts through different signaling pathways depending on the cell type and process^[7,8,28,30,61]^, a direct interaction between CNTN6 and the receptors of WNT and BMP/TGF-β pathways has not been established. Additionally, the modulation of these signaling pathways in *CNTN6*-mutant COs did not rescue all manifestations of the mutant phenotype, suggesting that CNTN6 may not directly influence the activity of these pathways. The most appealing molecular mechanism for CNTN6’s action is the Notch signaling pathway. Previous studies in rodents identified Cntn6 as a non-canonical ligand for the Notch1 receptor, triggering downstream pathway activation^[7,61]^. Our data extend these findings to humans, demonstrating interactions between CNTN6 and both NOTCH1 and NOTCH2 receptors. Additionally, *CNTN6* overexpression in NSCs activates Notch signaling. Furthermore, inhibition of Notch signaling in control COs phenocopies the key *CNTN6*-mutant defects, including impaired lumenization and disrupted PAX6 expression. Given the central role of Notch signaling both in neural morphogenesis and patterning, we propose that CNTN6 primarily exerts its effects through Notch signaling in human early cortical development, potentially via a non-cell-autonomous mechanism, as supported by our data.

Additionally, NICD1, released through the interaction between CNTN6 and NOTCH1, appears to regulate the intracellular trafficking of PAX6. Specifically, we observed aberrant cytoplasmic localization of PAX6 in some *CNTN6*-mutant cells, which was rescued upon *CNTN6* overexpression. A similar atypical subcellular localization has been described in Merkel cells in mice, where PAX6 translocated from the cytoplasm to the nucleus during differentiation^[62]^ and in the developing chick retina’s outer nuclear layer^[63]^. Notably, we observed PAX6 cytoplasmic localization only in NSCs with CNTN6 deficiency combined with 400-kb deletion. Given that simple loss of CNTN6 protein results in a reduction of PAX6, rather than complete suppression, normal nuclear localization in CNTN6-deficient NSCs may be compensated for by a complex PAX6 dose-dependent effect^[64,65]^ or its autoregulatory mechanisms^[66]^. However, we cannot exclude an additional unidentified effect of CNV.

Although there is no evidence of direct interaction between PAX6 and the NOTCH receptor, we hypothesized the existence of a molecular axis linking the PAX6 trafficking with the Notch pathway. One possible mechanism is that PAX6 is co-translocated with NICD from the cytoplasm to the nucleus. Our data indicate that NICD1 interacts with both PAX6(a) and PAX6(b) isoforms. It is well established that NICD forms complexes not only with the transcription factor CSL and the co-activator MAML but also with other proteins ^[67–69]^, including the transcription factor^[70]^. Although NICD is generally thought to bind its protein partners in the nucleus, emerging evidence suggests that such interactions may also occur in the cytoplasm^[71]^. Thus, our data do not exclude the existence of a link between PAX6 subcellular localization and the Notch pathway. However, the fundamental questions remain open: where is the complex between NICD and PAX6 formed? And is the interaction of these proteins CNTN6-dependent?

As noted above, the CNV in the *CNTN6* gene exhibits more pronounced pathological manifestations. Interestingly, most patients with neurodevelopmental disorders caused by mutations in *CNTN6* are predominantly carriers of CNVs^[1–3,72–75]^. CNV phenotypes are often unpredictable and can differ from those caused by point/indel mutations in the same gene^[76]^, due to varying molecular effects. We suggest that there are two most likely molecular effects of CNVs in *CNTN6*. The first is the “position effect,” which refers to changes in the regulation of gene expression. CNVs affect not only coding but also non-coding regions of genomes, such as non-coding RNAs^[77]^ or regulatory elements^[78–80]^, leading to the misregulation of neighboring genes. Therefore, we propose that some functional non-coding elements could be located within the deletion region of *CNTN6*, and their removal might disrupt the expression of adjacent genes. Notably, several genes (*CHL1, CNTN4, CRBN*) located near *CNTN6* are involved in cortical development, and mutations in these genes lead to various neurodevelopmental disorders^[81–86]^.

The second effect is the “two-hit” model, where CNVs create a sensitized genetic background that can influence the manifestation of other genetic changes^[87–89]^. Several case reports show that CNVs in *CNTN6* segregate with other pathological genetic variants^[90]^, supporting this model. Additionally, in some patients, *CNTN6* deletions or duplications are often a part of CNVs, affecting multiple genes simultaneously^[16,91–93]^. However, evaluating the contribution of an individual gene and determining its clinical significance in these cases is challenging. Nevertheless, the “two-hit” model may, to some extent, explain the incomplete penetrance and variable expressivity observed among CNV carriers in *CNTN6*^[1,17]^. Thus, CNVs in *CNTN6* probably may predispose to neurodevelopmental disorders by providing a genetic background for other pathogenic variants or creating a synergistic effect. However, more advanced and comprehensive diagnostics are needed to determine the precise pathogenetic mechanisms. In summary, our data uncover novel roles for the *CNTN6* locus in human cortical development, particularly in morphogenesis and tissue patterning. Additionally, our findings support and reinforce the contribution of CNVs to *CNTN6*-associated pathologies and suggest a synergetic effect between the CNVs at this locus and haploinsufficiency. Continued exploration of the underlying molecular mechanisms of neurodevelopmental disorders caused by variants in the *CNTN6* gene is essential to improving diagnostics and guiding potential therapeutic strategies.

### Experimental section

#### Human fetal tissue collection and preparation

The study was approved by the Ethics Committee of the Research Institute of Medical Genetics (protocol no. 10 from 15.02.2021). Written informed consent was obtained from the parents. A human fetus at stage GW8 was obtained following a medical pregnancy termination. Immediately after expulsion, the brain was removed and fixed for 24 h in 4% PFA (Sigma-Aldrich). For further histological studies, tissue was embedded in Tissue-Tek^®^ O.C.T. Compound (Sakura Finetek) and snap-frozen.

### Cell lines and culture conditions

#### HEK293T

HEK293T cells were cultured on 0.1% gelatin-coated (Sigma-Aldrich) culture ware in DMEM (Thermo Fisher Scientific), 10% FBS (Biosera), 2 mM L-glutamine, 1X MEM NEAA, and 1X Pen/Strep (all from Capricorn Scientific). For routine passaging, cells were treated with 0.25% trypsin-EDTA (BioinnLabs).

#### Human iPSC lines

The list of iPSC lines used in this study is presented in **Supplementary Table 1**, including source and RRIDs, where available. Several genetically modified iPSC lines were generated in-house and are not registered in public databases and the absence of RRIDs for these lines does not affect the validity or reproducibility of the conclusions. Cells were cultured on Corning^®^ Matrigel^®^ Matrix-coated (Corning) culture plates in mTeSR™1 complete medium (STEMCELL Technologies). For routine passaging every 5-7 days, cells were treated with StemPro™ Accutase™ Cell Dissociation Reagent (Thermo Fisher Scientific) for 5 min. ROCK inhibitor Y-27632 (STEMCELL Technologies) was added at a final concentration of 10 μM to increase the survival of iPSCs after dissociation. Cells were regularly tested and confirmed to be free of mycoplasma contamination by PCR.

### NSCs induction, cultivation, and terminal differentiation

Differentiation of iPSCs to NSCs was performed using the previously described protocol^[94]^ with minor modifications. Briefly, iPSCs were split and plated on the Corning^®^ Matrigel^®^ Matrix GFR (Corning) coated culture dish at a density of 100.000 cells/cm^2^. Differentiation was started when cells reached ∼100% confluence. The neural induction medium (50% DMEM/F12, 50% Neurobasal^TM^ medium (Thermo Fisher Scientific), 1X N-2 Supplement (Thermo Fisher Scientific), 1X B-27^TM^ Supplement (Thermo Fisher Scientific), 5 μg/ml insulin (Sigma-Aldrich), 2 mM L-glutamine, 1X MEM NEAA, 0.1 mM 2-mercaptoethanol, and 1X Pen/Strep) was refreshed daily. For the first 11 days, the medium was supplemented with 10 μM SB431542 and 2 μM Dorsomorphin (both from STEMCELL Technologies) to induce neural differentiation. On day 12, areas exhibiting neuroepithelial morphology were manually picked up and expanded on the Corning^®^ Matrigel^®^ Matrix GFR-coated dishes. Cells were subsequently passaged at a ratio of 1:2 to 1:3 and maintained in the neural induction medium with periodical cultivation in a neurospheres format. To induce terminal differentiation, NSCs were plated on the Corning^®^ Matrigel^®^ Matrix GFR-coated plate at a density of 25.000 cells/сm^2^ in neural induction medium supplemented with 10μM Y-27632. The next day, the medium was changed to BrainPhys™ Neuronal Medium supplemented with 1X NeuroCult™ SM1 Neuronal Supplement, 1X N2 Supplement-A, 20 ng/ml Human Recombinant BDNF, 20 ng/ml Human Recombinant GDNF (all from STEMCELL Technologies), 10 μM Forskolin (Sigma-Aldrich), 200 nM ascorbic acid (Wako), and 1X Pen/Strep. The medium was changed every 3-4 days for 3 weeks.

### Cerebral organoid culture

The generation of COs was performed according to the classic Lancaster’s protocol^[95]^ with minor modifications. Briefly, iPSCs were dissociated into a single-cell suspension using StemPro™ Accutase™ Cell Dissociation Reagent and stained with trypan blue solution (Corning) to identify dead cells. The number of live cells was counted with a hemocytometer, and 3.000.000 cells were plated on an AggreWell™800 microwell culture plate (STEMCELL Technologies) prepared according to the manufacturer’s protocol. Y-27632 and bFGF were added to EB medium (80% DMEM/F12, 20% KnockOut™ Serum Replacement, 2 mM L-glutamine, 1X MEM NEAA, 0.1 mM 2-mercaptoethanol, and 1X Pen/Strep) to a final concentration of 10 μM and 4 ng/ml, respectively. Half of the medium was replaced the next day with fresh EB medium. On day 2, EBs were dislodged from the AggreWell™800 plate and transferred to culture plastic pre-treated with Anti-Adherence Rinsing Solution (STEMCELL Technologies). The medium was changed every day. bFGF was removed from the medium on day 5 for the next 48 h. Then, the medium was completely replaced with neural induction medium (96% DMEM/F12, 1X N-2 Supplement, 2 mM L-glutamine, 1X MEM NEAA, 0.1 mM 2-mercaptoethanol, 1X Pen/Strep, and 1 μg/ml heparin (Sigma-Aldrich)) and refreshed every two days. Afterwards, EBs were embedded in Corning^®^ Matrigel^®^ Matrix GFR and cultivated until day 15 in cerebral organoid medium VitA-(50% DMEM/F12, 50% Neurobasal^TM^ medium, 0.5X N-2 Supplement, 1X B-27^TM^ Supplement w/o vitamin A (Thermo Fisher Scientific), 2 mM L-glutamine, 0.5X MEM NEAA, 0.1 mM 2-mercaptoethanol, 1X Pen/Strep, and 2.5 μg/ml insulin). On day 15, organoids were removed from the matrigel droplets, and the medium was replaced with cerebral organoid medium VitA+ (50% DMEM/F12, 50% Neurobasal^TM^ medium, 0.5X N-2 Supplement, 2X B-27^TM^ Supplement, 2 mM L-glutamine, 1X MEM NEAA, 0.1 mM 2-mercaptoethanol, 1X Pen/Strep, 2.5 μg/ml insulin, 0.8 mM ascorbic acid, and 20 mM HEPES (Capricorn Scientific)) and started agitation (90 rpm). Refreshing of the medium was performed every 3-4 days until the end of the experiments. An orbital shaker (Celltron, Infors) was used at 90 rpm for 10 cm dishes. For the generation of chimeric COs, iPSCs from lines iTAF1-36 and iTAF3-17 C9-8-GFP were mixed at a ratio of 60:1, respectively, before plating on an AggreWell™800 microwell plate. COs were treated with 2 μg/ml Doxycycline for rescue phenotype experiments. Control COs were treated with PBS. In BMP/TGF-β and Notch pathway inhibition experiments, small molecules SB431542, Dorsomorphin, and DAPT (Cell Signaling Technology) were used at concentrations of 10 μM, 2 μM, and 10 μM, respectively. Control COs were treated with DMSO (BioFroxx).

### Genome editing with CRISPR/Cas9

#### Generation of iPSC lines with endogenous tagging of *CNTN6* exon 2

Human iPSC lines iTAF3-17 and iTAF1-36 were used as the parental lines to generate *CNTN6* knockout cell lines. CRISPR/Cas9-mediated genome editing was applied to introduce indel mutations through non-homologous end joining (NHEJ) and to insert a stop-cassette into exon 2 of the *CNTN6* gene with ssODN donor through homology-directed repair (HDR). Guide RNAs were cloned into a gRNA cloning vector (Addgene, #41824). The stop cassette flanked by homology arm sequences was designed to contain stop codons in each open reading frame and HindIII restriction site to simplify genotyping. Transfection of plasmid vectors coding for gRNA and Cas9 protein (Addgene, #127762 or #62988) alone (for NHEJ) or with adding ssODN donor (for HDR) was performed either with Neon™ Transfection System (Thermo Fisher Scientific) using the settings 1100 V, 20 ms, 1 pulse, according to the manufacturer’s instructions or standard transfection with Lipofectamine^®^ 3000 (Thermo Fisher Scientific). The day after transfection, cells were plated on a murine embryonic fibroblast feeder layer and cultivated in human iPSC medium (80% DMEM/F12, 20% KnockOut™ Serum Replacement (both from Thermo Fisher Scientific), 2 mM L-glutamine, 1X MEM NEAA, 1X Pen/Strep, 0.1 mM 2-mercaptoethanol (Helicon), and 10 ng/ml bFGF (Thermo Fisher Scientific)), for selection cells were treated with of 0.35 μg/ml puromycin (Thermo Fisher Scientific) or 400 μg/ml G418 (Enzo Life Sciences) for 5-7 days. Next, the formed colonies were handpicked, expanded, and analyzed using PCR, restriction, and Sanger sequencing. Oligonucleotide sequences for cloning and primers are disclosed in **Supplementary Table 8.**

### Generation of iPSC lines with modified *AAVS1* locus

For rescue assay, the coding region of the *CNTN6* gene was amplified with specific primers from human cDNA using PCR and cloned into an AAVS1 donor vector for inducible gene expression (Addgene, #52343). *AAVS1* targeting of iTAF3-17 C9-8 cells was performed by nucleofection using the Neon™ Transfection System, as described previously. After transfection, cells were selected with puromycin (0.35 μg/ml) for 7 days, and the single clones were picked based on their GFP expression after induction with 2 μg/ml Doxycycline (Sigma-Aldrich), then expanded and analyzed by PCR with previously published primers^[96]^.

### Fixation and cryosectioning

COs were fixed in 4% PFA for 1 h on the roller shaker at RT. Samples were washed thrice with PBS for 20 min each wash. For dehydration, COs were incubated overnight in a 15% sucrose (Sigma-Aldrich) solution at +4°C. The next day, the solution was changed to a 30% sucrose solution. Dehydrated COs were embedded in a gelatin/sucrose solution (10%/7.5%, respectively), snap-frozen, and stored at -80°C. Cryosectioning was performed using a cryostat Microm HM550 or CryoStar NX70 (both from Thermo Fisher Scientific). Sections with a thickness of 30-40 μm were attached on Superfrost PLUS slides (Epredia), dried overnight at RT, and kept at -20°C.

Coverslips with cells were fixed in 4% PFA for 15 min at RT. Then, coverslips were washed in PBS thrice and kept in PBS at 4°C. For extended storage, sodium azide (Sigma-Aldrich) was added to a final concentration of 0.03%.

### Immunohistochemistry

Slides were washed twice with PBS and incubated with a blocking buffer (2% BSA (Sigma-Aldrich), 5% FBS (Thermo Fisher Scientific), and 0.2% Triton (Medigene) and primary antibodies overnight at RT on an orbital shaker (50 rpm). Information about antibodies is presented in **Supplementary Table 9.** Next, slides were washed thrice with PBS and incubated with secondary antibodies and DNA stains (DAPI or Hoechst 33258; both from Sigma-Aldrich). Slides were washed with PBS three times, thoroughly dried, and covered with ProLong™ Diamond Antifade Mountant (Thermo Fisher Scientific) or Mounting medium (Servicebio Technology). Immunostaining of coverslips was performed using the same procedure.

### Karyotyping

Karyotype analysis of iPSCs was performed according to the previously described cytogenetic protocol^[97]^ with minor modifications. Briefly, cells at ∼80% confluency were treated with colcemid (Capricorn Scientific GmbH) at a final concentration of 50 ng/ml for 3 h before fixation. Cells were incubated with 0.05% trypsin-EDTA for 5 min at 37°C. Then, a hypotonic solution (0.38 M KCl) was added to the cells and incubated for 20 min at 37°C. Next, the cells were prefixed by adding 2-3 drops of the fixative (3:1 methanol/acetic acid, respectively), mechanically removed from the dish, and centrifuged at 1300 rpm for 6 min. The cell pellet was washed twice with 1.5 ml cold fixative and incubated on ice for 20 and 10 min between washes. The cell pellet was resuspended in 300 µl of cold fixative, and 100 µl of cell suspension was applied onto prechilled slides, incubated on ice for 3-5 min, and thoroughly dried under hot air. The preparations were stained with 0.1 µg/ml DAPI for 3 min, washed in 2X SSC buffer and distilled water, and covered with an antifade mountant (1% DABCO (Sigma-Aldrich), 90% glycerol (MP Biomedicals), 1X PBS). The images of metaphase plates were recorded and processed using Ikaros software (MetaSystems). At least 50 metaphases were analyzed for each iPSC line (450-band resolution). Chromosomes were described using ISCN2016^[98]^.

### Microscopic analysis and imaging processing

EBs were acquired with a Zeiss Axio Observer Z1 in brightfield mode. Fluorescent images of live cells and COs were generated with a ZOE Fluorescent Cell Imager (Bio-Rad). Imaging of fixed cells and CO sections was performed using LSM 510 META, LSM 780 NLO (both from Zeiss), and Fluoview FV3000 (Olympus) confocal microscopes. Image processing was performed in ImageJ software^[99]^. Cross-sectional areas of EBs, VZ-like tissue area and thickness, PAX6+ tissue area, and ZO-1+ apical membrane length were measured with the freehand selection tool. The number of PH3+ and total cells was measured by the multi-point tool. Mitotic spindle angles were measured with the angle tool. Inflection points in COs were measured according to the previous publication^[100]^.

### Bulk RNA-seq

COs were washed in ice-cold PBS and lysed in TRI Reagent^®^ (Sigma-Aldrich). Total RNA was extracted using the Direct-zol™ RNA Miniprep Plus Kit (Zymo Research), following the manufacturer’s instructions. RNA sequencing was performed by BGI Genomics. Each sample accounted for about 62 million reads.

### Computational methods

#### RNA-seq data processing

The quality of the raw sequencing reads was assessed using FastQC^[101]^. Subsequent read filtering was conducted with FASTP^[102]^, applying default parameters. Transcript quantification was carried out using Salmon^[103]^, with transcript annotation provided by GENCODE GRCh38.p13. Differential gene expression analysis was performed using DESeq2^[104]^. To generate a list of common DEGs, we overlapped DEGs from the comparisons between Control #1 versus *CNTN6^Δ/mut^* and Control #1 versus *CNTN6^mut/mut^* groups. Gene ontology analysis for all DEG groups was conducted using Metascape^[105]^.

#### Analysis of Notch target genes enrichment

We conducted an enrichment analysis of the DEGs for Notch target genes identified in previously published studies^[31,32]^. Initially, we identified human orthologs corresponding to the mouse genes reported in these studies. Subsequently, we assessed the enrichment of Notch target genes within the DEGs using the hypergeometric distribution, as implemented in the scipy.stats Python library. The total number of human genes (61272) was used as the total number of objects for this calculation.

### DNA cloning

Genetic constructs used for overexpression of specific genes were based on pCAGGS-mCherry (Addgene Plasmid #41583), into which 3xHA, 1xFLAG or eGFP tags were inserted at the 3’-end for Co-IP experiments. Human gene coding sequences (CDSs) were amplified with specific primers (**Supplementary Table 8**) from COs and HEK293 cDNA with Q5 polymerase (NEB) and placed into respective vectors either via EcoRI/AgeI cloning sites or by Gibson assembly (NEBuilder® HiFi DNA Assembly Master Mix). The regions that were cloned: 1-723 aa DLL1 (NM_005618.4); 1-999 aa CNTN6 (NM_014461.4); 1-1796 aa NECD1, 1753-2555 aa NICD1 (NM_017617.5); 1-1677 aa NECD2, 1928-2471 aa NICD2 (NM_024408); 1-422 PAX6a (NM_001368887); 1-436 aa PAX6b isoform (NM_001368892). A list of the genetic constructs is provided in **Supplementary Table 10**. All sequences and vectors are available upon request.

### Plasmid transfection

NSCs were transfected using GenJect™-39 (Molecta) according to the manufacturer’s recommendations. Briefly, cells were seeded at a density of 100.000 cells/cm^2^ on a 24-well plate. The following day, cells were transfected with a mix of plasmid DNA and GenJect™-39 reagent at a ratio of 1:1. The total amount of plasmid DNA used was 1.5 μg per well. Cells were incubated for 6 h with the mix, after which the medium was replaced with fresh NSC medium. Transfected NSCs were harvested for analysis 48 h post-transfection.

HEK293T cells were transfected using polyethylenimine (PEI) (Sigma Aldrich). Briefly, cells were seeded at 70.000 cells/cm^2^. For transfection, 2 µg of plasmid DNA was mixed with PEI at a ratio of 2 µg per 1 µg DNA. The transfection complex was incubated at room temperature for 20 min. The mix of plasmid DNA and PEI was added to the cells and incubated overnight. The next day, the medium was changed, and the cells were harvested for analysis 48-72 h post-transfection.

### Protein sample preparation

Сells or COs were harvested and lysed in IP Lysis Buffer (25 mM Tris-HCl pH 7.4, 150 mM NaCl, 1 mM EDTA, 1% NP-40, 5% glycerol) supplemented with a protease inhibitor cocktail (5892970001, Roche), PhosSTOP™ (4906845001, Roche) and 5 mM NaF (S2002, Sigma-Aldrich). Samples were incubated for 15 min on ice and sonicated for 10 s at 35% power. After centrifugation for 20 min at 4°C, supernatants were collected into fresh tubes. The protein concentration of samples was measured using Pierce™ BCA Protein Assay Kit (23225, Thermo Fisher Scientific) according to producer recommendations. Colorimetric detection was performed with the CLARIOstar^®^ Plus microplate reader (BMG LABTECH).

### Co-immunoprecipitation

200 µl of protein samples with a final concentration of 1 mg/ml were precleared by incubation with Protein A or G magnetic beads (Cell Signaling Technology or Sileks) for 30 min at 4°C. Pre-cleared samples were incubated with primary antibodies or IgG isotype controls (**Supplementary Table 9**) with rotation overnight at 4°C. Then, immune complexes were precipitated on magnetic beads with rotation for 1 h at RT. After washing with IP Lysis Buffer, samples were eluted in 3X SDS Buffer (180 mM Tris-HCl pH 6.8, 3% SDS, 30% glycerol, 0.01 % bromophenol blue, 10% 2-mercaptoethanol, 100 mM DTT) by boiling at 95°C for 5 min.

### Western Blotting

Protein samples were loaded with 20 μg per lane. Proteins were separated under denaturing conditions using SDS-PAGE. Protein transfer was performed on the PVDF membrane (0.42 μm pore size, Bio-Rad) at 90 V for 2 h. Membranes were blocked in 5% milk (Cell Signaling or Neofroxx) for 1 h at RT. Staining with primary antibodies was performed overnight on a roller shaker in 5% non-fat milk in TBST. Membranes were washed three times with TBST for 30 min following incubation with HRP-conjugated secondary antibodies. Detection was performed with Clarity™ Western ECL Substrate (Bio-Rad) with iBright Imaging Systems (Thermo Fisher Scientific). For the detection of low-expressing protein, SuperSignal™ West Atto Ultimate Sensitivity Substrate was used (Thermo Fisher Scientific).

Antibody stripping was performed according to a previous publication^[106]^. Briefly, membranes were washed twice in TBST and incubated twice with stripping buffer (6 M GnHCl, 0.2% Nonidet P-40, 0.1 M 2-mercaptoethanol, and 20 mM Tris–HCl, pH 7.5) for 10-15 min at RT on an orbital shaker. Next, they were washed three times with TBST for 10 min each, blocked in 5% non-fat milk in TBST, and reprobed with primary antibodies.

For protein quantification, images of membranes were analyzed with iBright™ Analysis Software (Thermo Fisher Scientific). Local Background Correction Volume was counted for each protein. The normalization factor for each lane was calculated as a ratio between the brightest and other housekeeping bands. The quantification of target proteins was normalized to GAPDH, and fold change was counted for at least two independent experiments.

## Acknowledgments

We are grateful to all families for their participation in the study. We also extend our special gratitude to Alexey Pindyurin, Anna Malashicheva, Ingo Bormuth, and Olga Bormuth for their insightful discussions and constructive feedback. This study was funded by the Russian Foundation for Basic Research (grant no. 19-29-04067), Russian Science Foundation (grant no. 14-15-00772), and the Ministry of Science and Higher Education of the Russian Federation (state project no. FWNR-2022-0019 and state contract no. 075-15-2021-1063). NGS data analysis was performed on the nodes of the Novosibirsk State University high-throughput computational cluster, supported by the Ministry of Science and Higher Education of the Russian Federation (grant no. FSUS-2024-0018). We also acknowledge the Multiple-Access Center for Microscopy of Biological Subjects, ICG SB RAS, and personally to Sergei I. Baiborodin for technical assistance with the microscopic equipment. Additionally, we thank the Collective Center of ICG SB RAS “Collection of Pluripotent Human and Mammalian Cell Cultures for Biological and Biomedical Research” for providing access to favorable conditions for culturing human iPSCs and COs. Schematics were generated using BioRender (https://app.biorender.com/).

## Statistical analysis

Statistical analysis was performed using BioRender (https://app.biorender.com/). The raw values are provided in **Source Data 1**. Detailed information for each graph, including sample size, statistical tests, and corresponding P-values, are indicated in the figure legends. The Shapiro-Wilk test was used to assess data normality. For comparisons between two groups, the Mann-Whitney test was applied, while the Kruskal-Wallis test with post-hoc Dunn’s analysis was used for multiple group comparisons.

## Data availability

The raw sequencing RNA-seq data have been submitted to the NCBI database with bioproject accession number PRJNA1165993.

## Conflict of Interest

The authors declare no conflict of interest.

---

**Figure S1.**
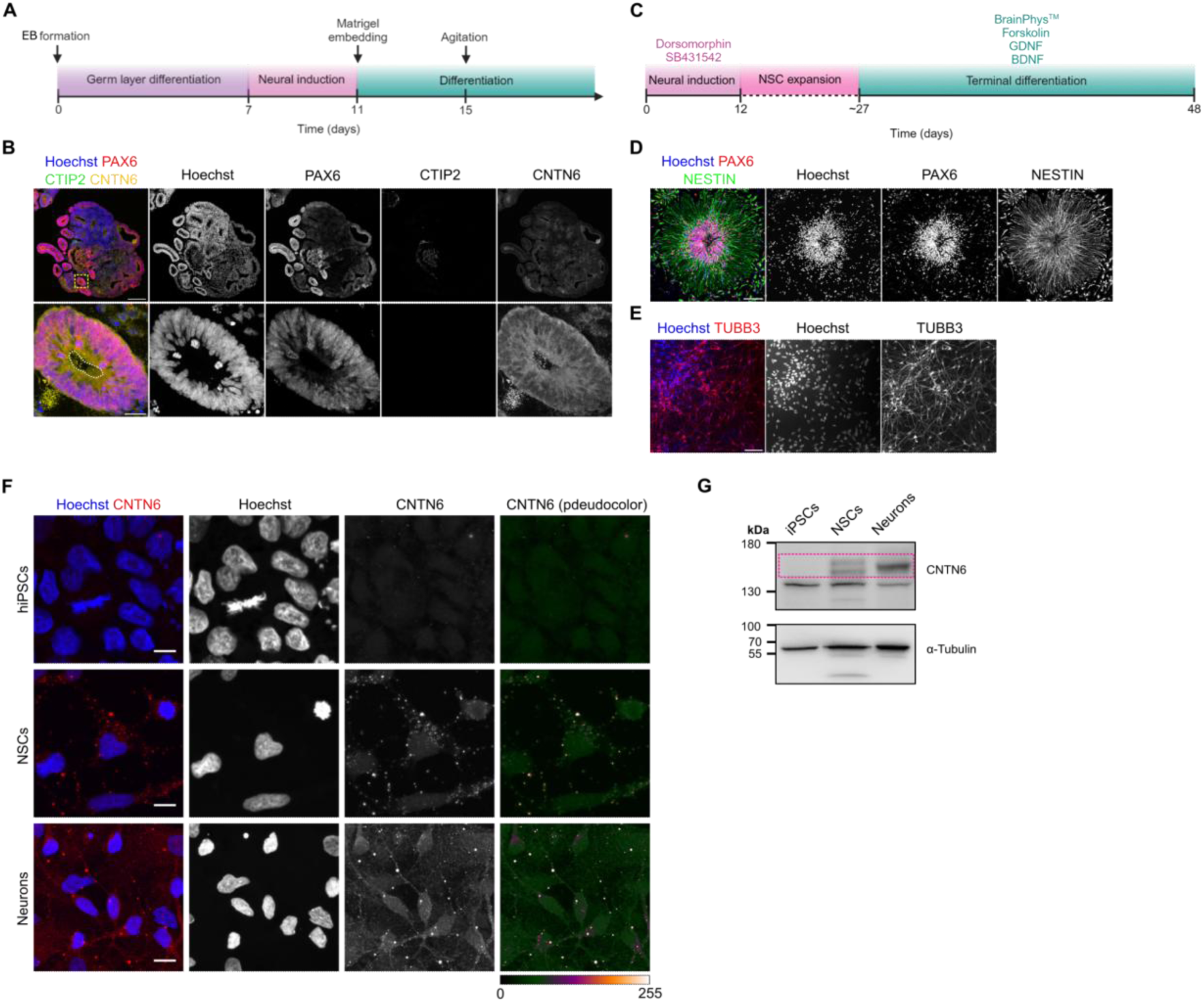
Spatiotemporal expression of CNTN6 during the early stages of human cerebral cortex development. **A** Schematic of the protocol for CO generation. **B** Immunohistochemical analysis of CNTN6 expression in day 20 COs. Separate channels are presented in grayscale. CTIP2 and PAX6 mark deep-layer neurons and radial glial cells, respectively. The white dotted line labels the apical surface, yellow dotted square labels the ROI. Scale bar: 250 μm (upper panel) and 50 μm (bottom panel). **C** Schematic of the protocol for the generation of NSCs and neurons from iPSC by dual SMAD inhibition. **D, E** Immunohistochemical analysis of the NSCs and neurons derived from iPSCs. Separated channels are presented in grayscale. NESTIN and PAX6 label NSCs (**D**), and TUBB3 marks neurons **(E)**. Scale bar: 100 μm (**D**) and 50 μm (**E**). **F** Immunohistochemical analysis of CNTN6 expression in iPSCs, NSCs, and neurons. Separated channels are presented in grayscale, and a pseudocolor representation of the signal distribution is presented for the CNTN6 channel. iPSCs were used as a negative control. Scale bar: 10 μm. **G** Representative western blot results of CNTN6 protein expression in iPSCs, NSCs, and neurons. Pink dotted rectangle labels bands corresponding to CNTN6.

**Figure S2.**
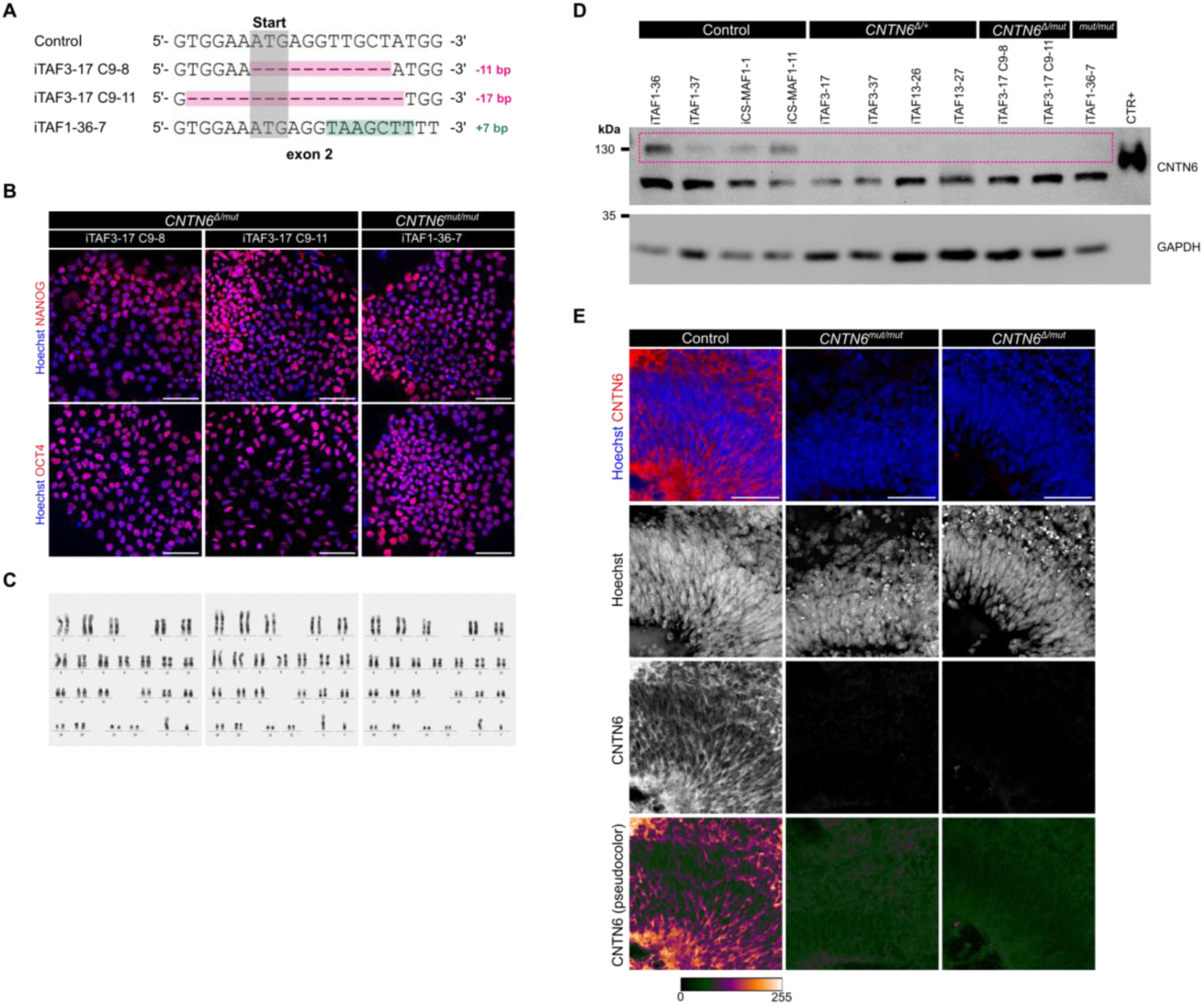
Generation and characterization of *CNTN6* mutant iPSC lines. **A** Alignment of DNA sequences demonstrates CRISPR/Cas9-mediated gene editing of exon 2 in the *CNTN6* gene in mutant iPSC lines. The gray rectangle highlights the start-codon, the pink rectangle indicates deletion, green – the insertion. **B,C** Characterisation of *CNTN6* mutant iPSC lines. Representative images of immunohistochemical analysis **(B)** of iPSC lines for pluripotent markers expression NANOG and OCT4 (both red). Scale bar: 100 μm. **(C)** Karyograms reveal normal karyotypes in iPSC lines. **D** Representative western blot results of CNTN6 protein in iPSC-derived COs at 90 days of differentiation, confirming *CNTN6* gene knockout. The dotted pink rectangle indicates the CNTN6 protein (∼130 kDa). The lowest band row is nonspecific staining. GAPDH protein is used as a housekeeping control. CTR+ – recombinant CNTN6 is used as a positive control. **E** Immunohistochemical analysis of CNTN6 expression on day 90 CO, confirming *CNTN6* gene knockout. Separated channels are presented in grayscale. In addition, a pseudocolor representation of the signal distribution is presented for the CNTN6 channel. Scale bar: 50 μm

**Figure S3.**
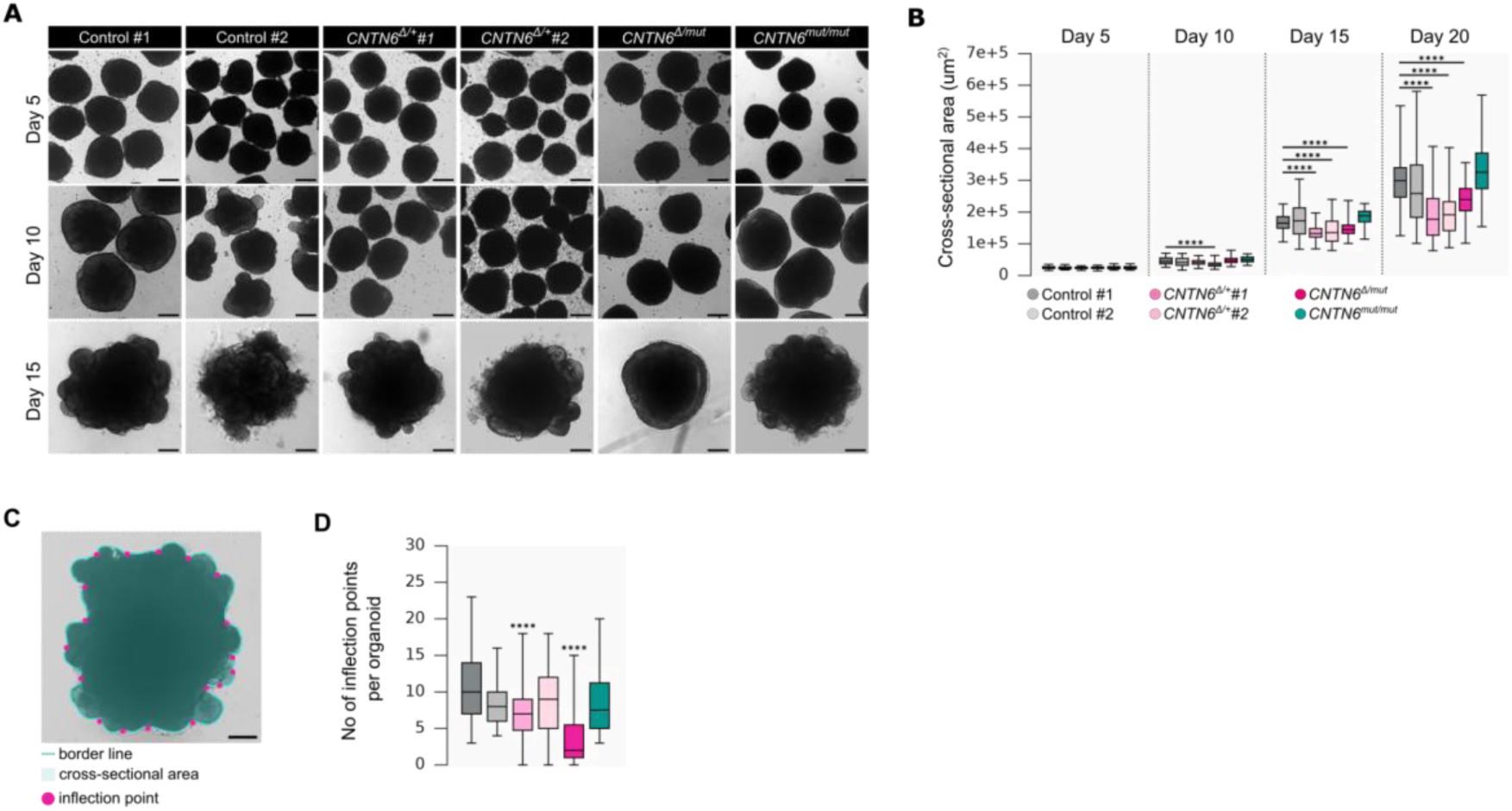
CNV and knockouts in the *CNTN6* gene lead to disruption of VZ-like structure formation during CO differentiation. **A** Representative brightfield images of COs on days 5, 10, and 15. Scale bar: 100 μm. **B** Quantification of the cross-sectional area of COs. Box plots represent median and quartiles with minimum and maximum values (n>40 for each group on any day; ****P < 0.0001; Kruskal–Wallis test with post-hoc Dunn’s test). **C** Schematic of inflection point measurements. Scale bar: 100 μm. **D** Quantification of the inflection points per organoid at day 20. Box plots represent median and quartiles with minimum and maximum values (n>32 for each group; ****P < 0.0001; Kruskal–Wallis test with post-hoc Dunn’s test).

**Figure S4.**
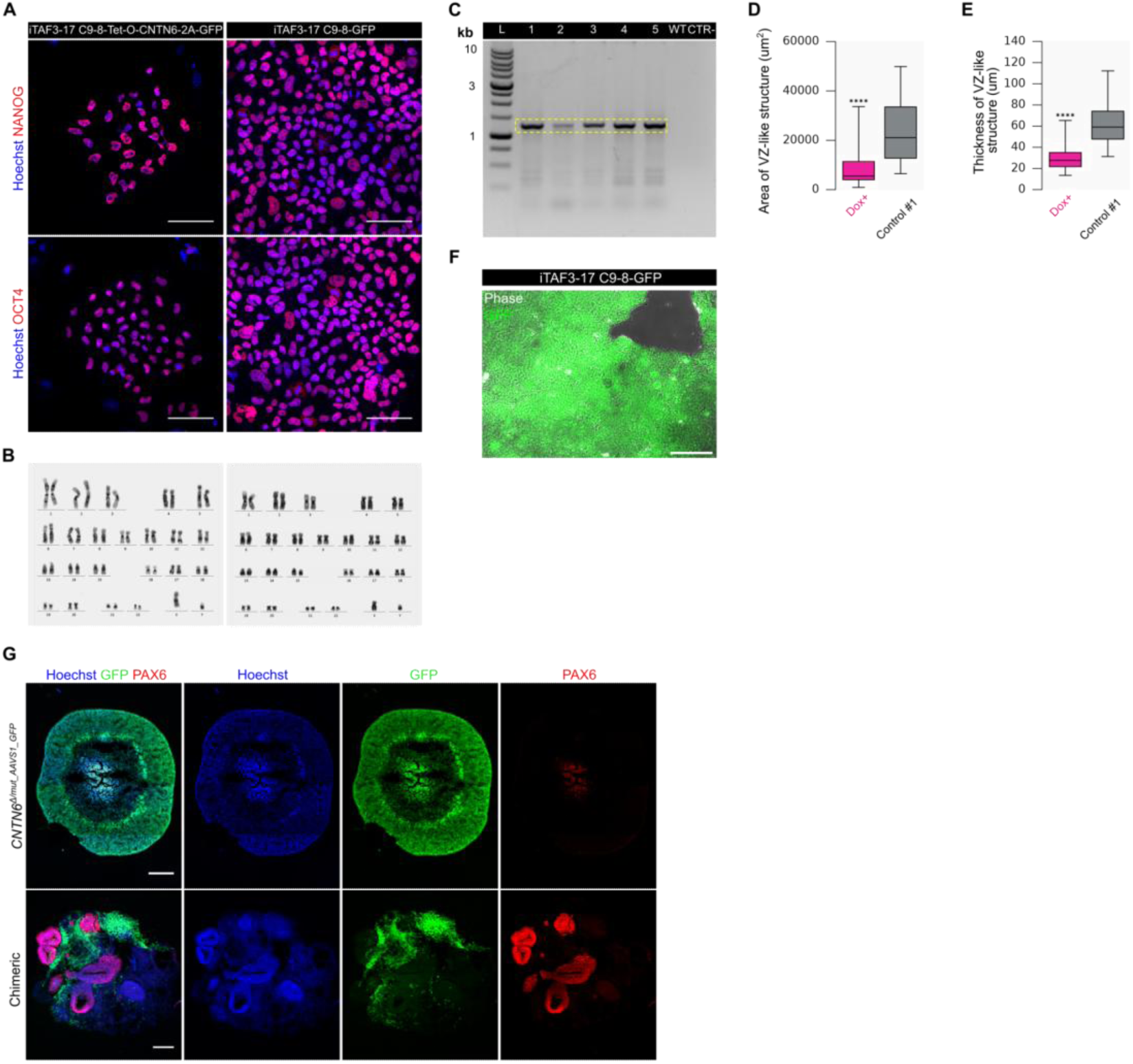
Partial rescue of phenotype in *CNTN6^Δ/mut^* COs by *CNTN6* overexpression and chimeric organoid approach. **A, B** Characterisation of *CNTN6^Δ/mut_AAVS1-CNTN6-GFP^ CNTN6^Δ/mut_AAVS1-GFP^* iPSC lines with modified *AAVS1* locus. Representative images of immunohistochemical analysis **(A)** of iPSC lines for the expression of pluripotent markers, NANOG and OCT4 (both red). Scale bar: 100 μm. **B** Karyograms reveal normal karyotypes in iPSC lines. **C** Representative PCR genotyping of iPSC clones. The yellow dotted rectangle indicates bands with the corresponding insert into the *AAVS1 l*ocus (L – ladder, WT – original iPSC line, CTR-– negative PCR control). **D, E** Quantitative analysis of areas and thickness of VZ-like structures in *CNTN6^Δ/mut^*COs with CNTN6 overexpression and Control #1 COs. Box plots represent median and quartiles with minimum and maximum values (Control #1 n=58, Dox+ n=50; ****P < 0.0001; Mann-Whitney U test). **F** Representative combined brightfield and fluorescent image of *CNTN6^Δ/mut^* iPSCs with GFP constitutive overexpression. Scale bar: 250 μm. **G** Immunohistochemical analysis of PAX6 expression in chimeric and *CNTN6^Δ/mut^* COs at day 20. Scale bar: 200 μm.

**Figure S5.**
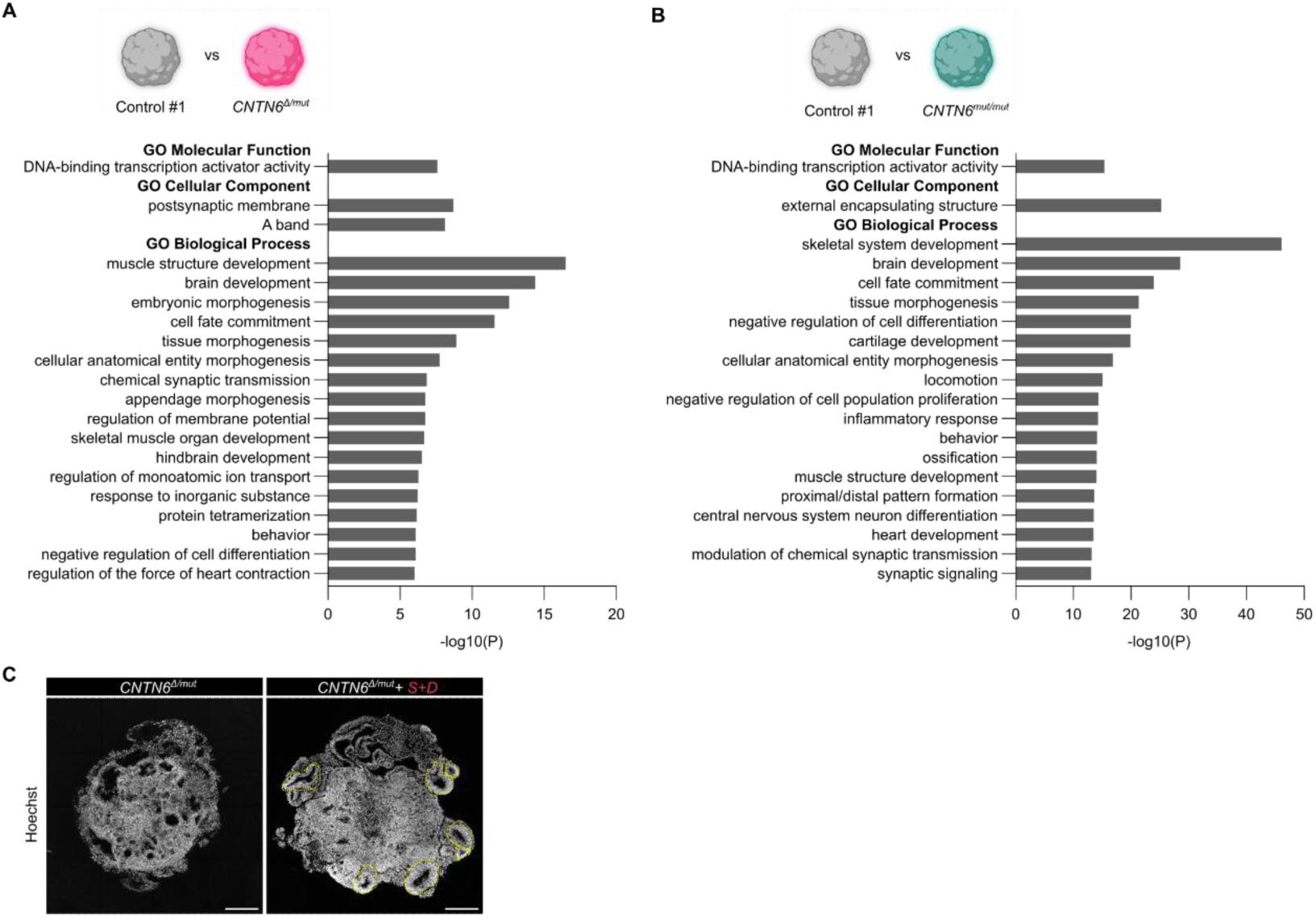
Comparative gene expression analysis of COs with deletion and knockouts in the *CNTN6* locus. **A,B** GO analysis of DEGs using Metascape (https://metascape.org/) identified from CO RNA-seq is shown. Metascape uses the hypergeometric test for enrichment P-value calculation. The bar chart represents -Log10 P-values for the 20 top-ranked GO terms. **C** Representative fluorescent images of the entire organization of *CNTN6^Δ/mut^* COs on day 20 under inhibition of the BMP/TGFβ signaling pathway. The yellow dotted line labels VZ-like structures. Scale bar: 250 μm.

## Notes

### Competing Interest Statement

The authors have declared no competing interest.

