## Supplementary material for "A novel role for the *CNTN6* locus in lumenization and radial glial cell fate determination during early human cortical development revealed in cerebral organoids": Source Data 2

### Source Data related to Figure 1c

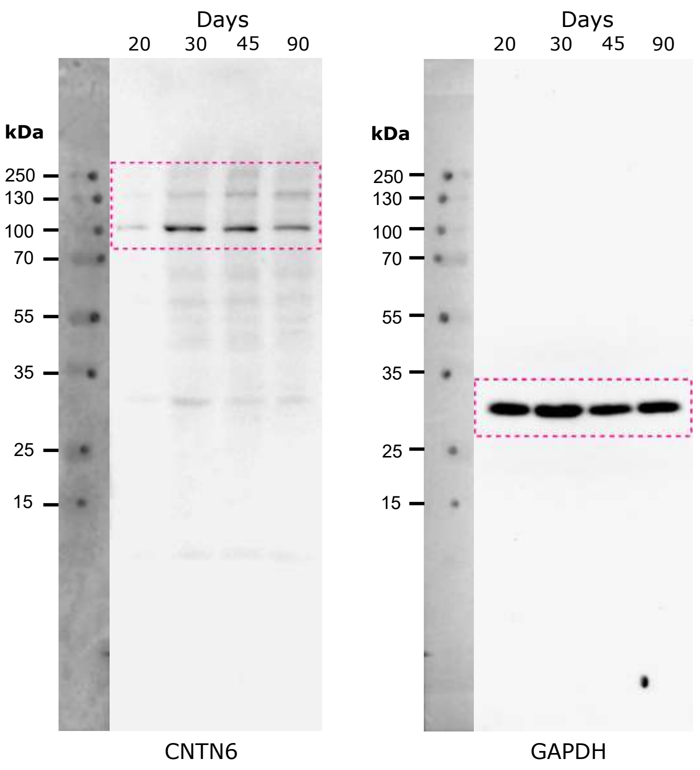


Pink dotted line marks the region of interest that has been cropped and represented in Figure 1c.

Source Data related to Extended Data Figure 1g

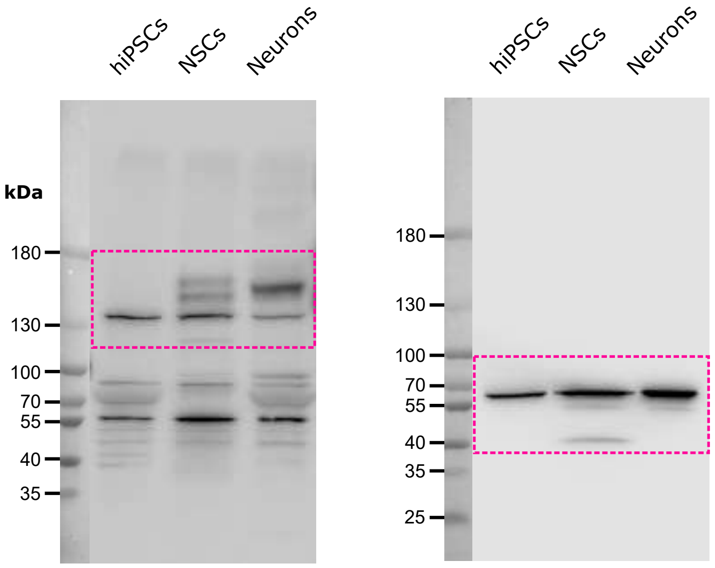


Pink dotted line marks the region of interest that has been cropped and represented in Extended Data Figure 1g.

### Source Data related to Extended Data Figure 2d

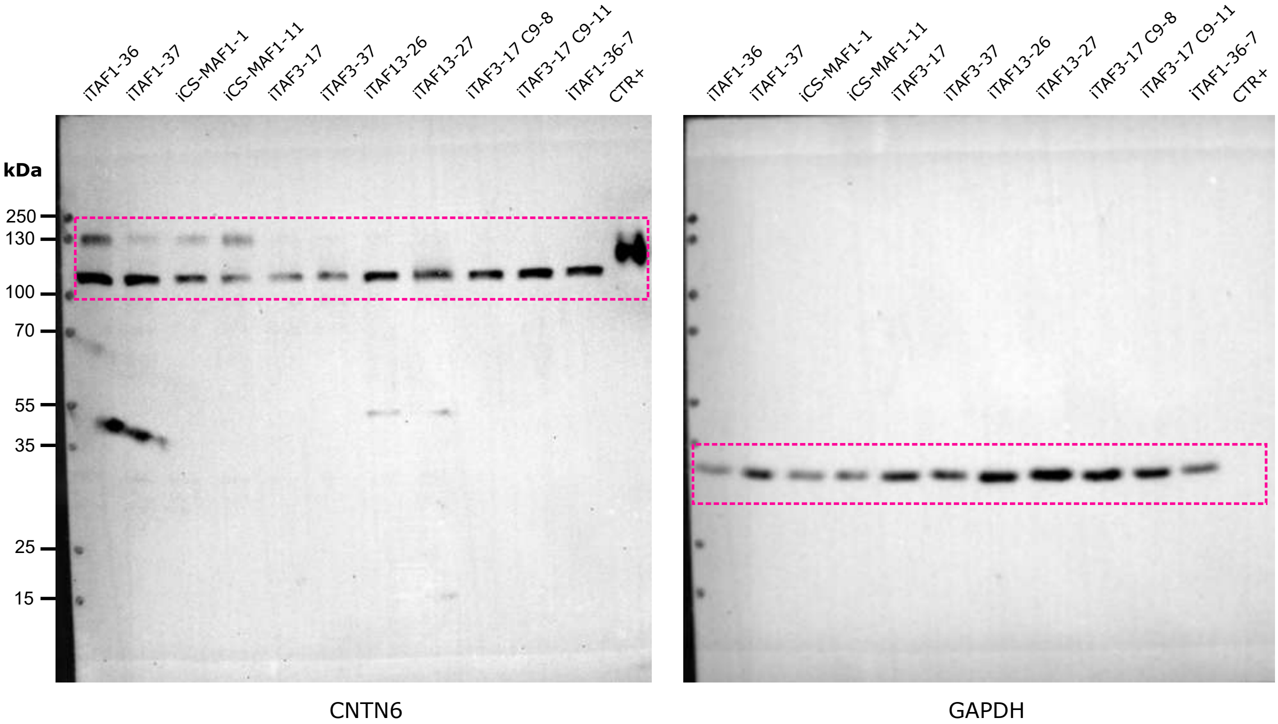


Pink dotted line marks the region of interest that has been cropped and represented in Extended Data Figure 2d.

Source Data related to Figure 4d

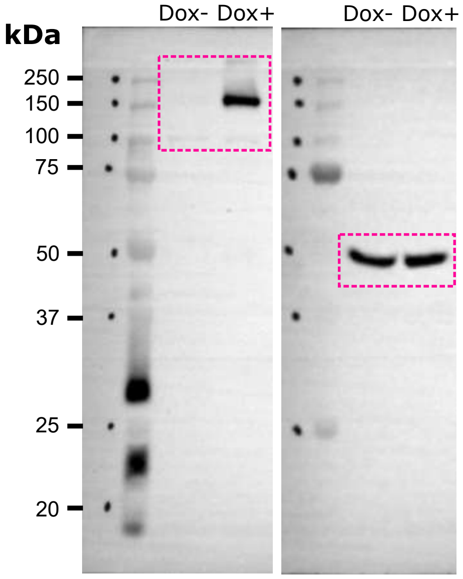


Pink dotted line marks the region of interest that has been cropped and represented in Figure 4d.

### Source Data related to Figure 6b

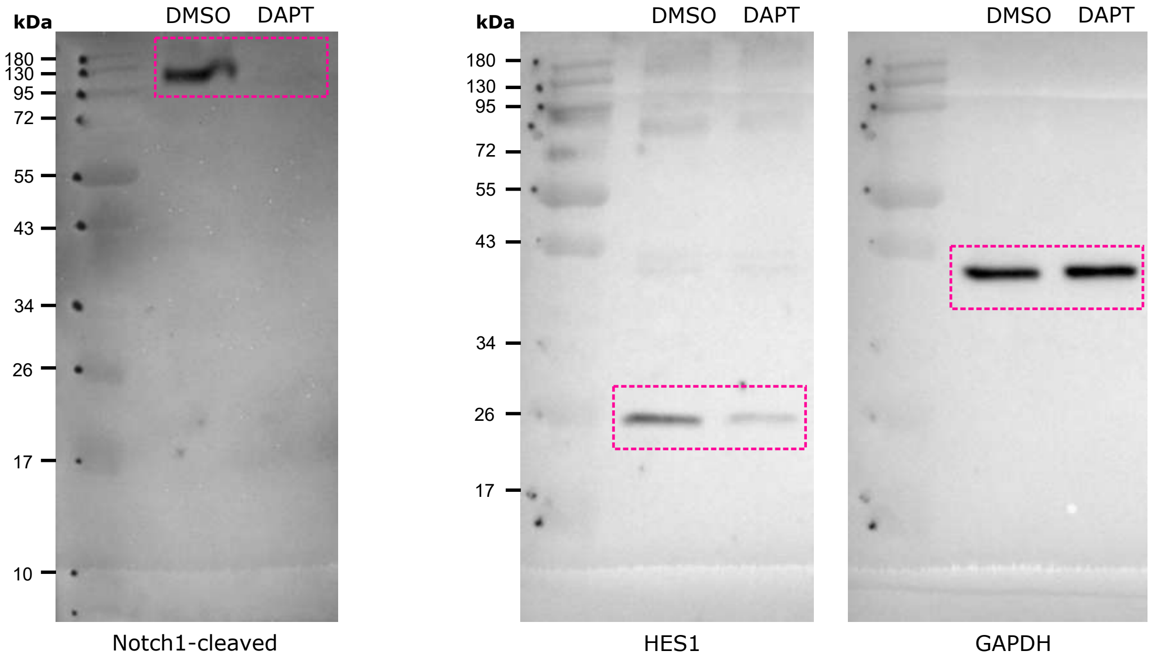


Pink dotted line marks the region of interest that has been cropped and represented in Figure 6b.

### Source Data related to Figure 7

#### Figure 7a

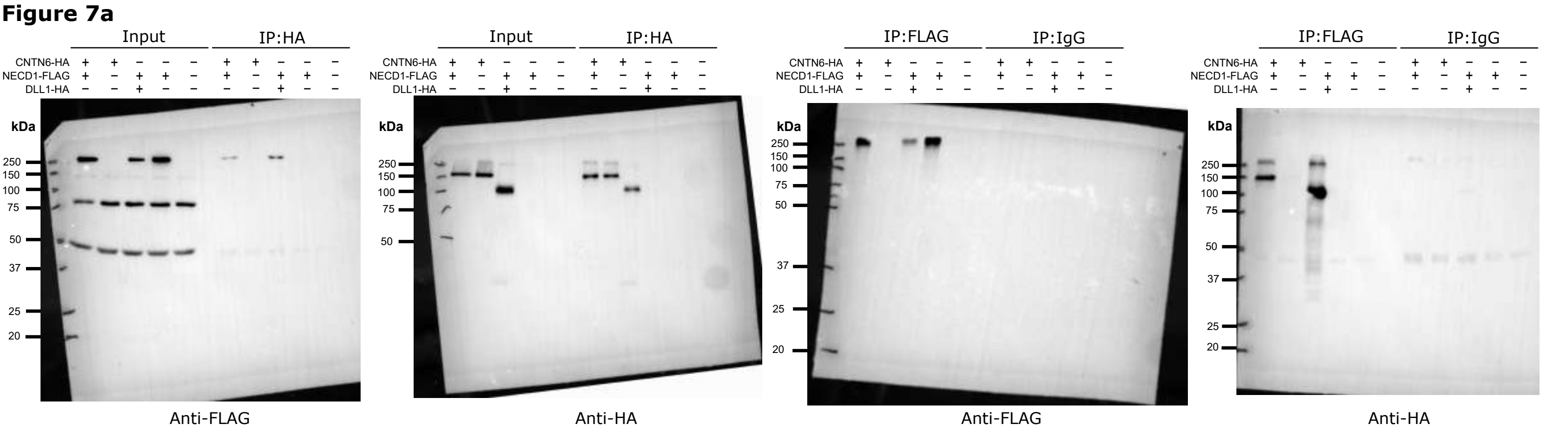


#### Figure 7b

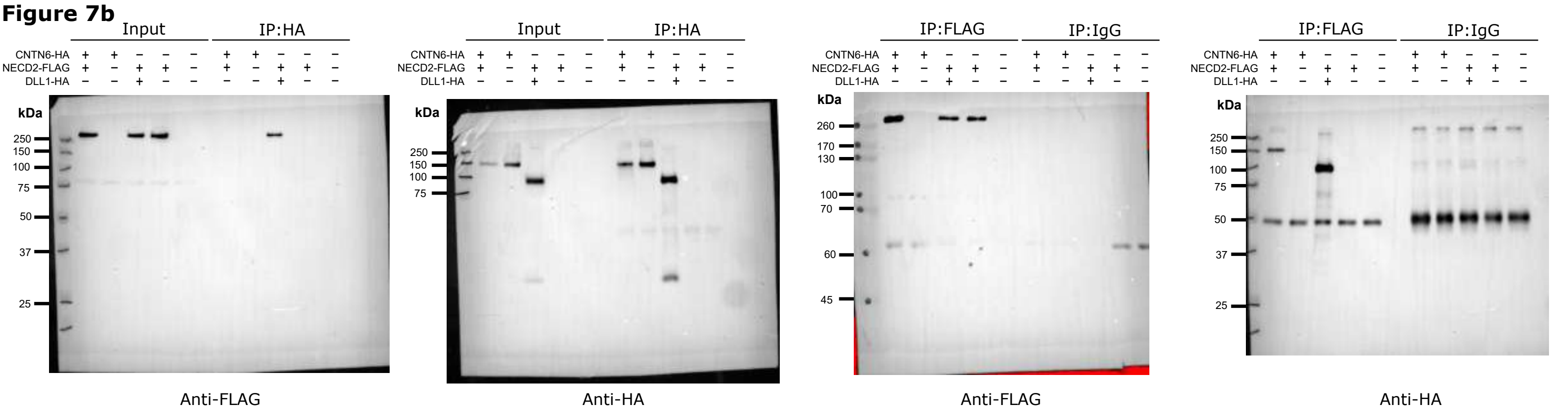


#### Figure 7c

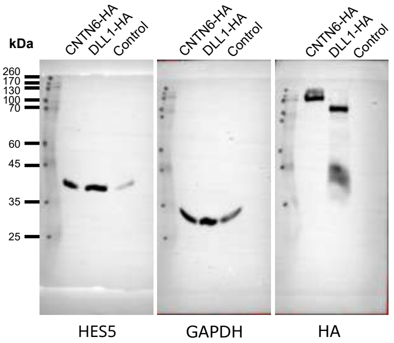


#### Figure 7f

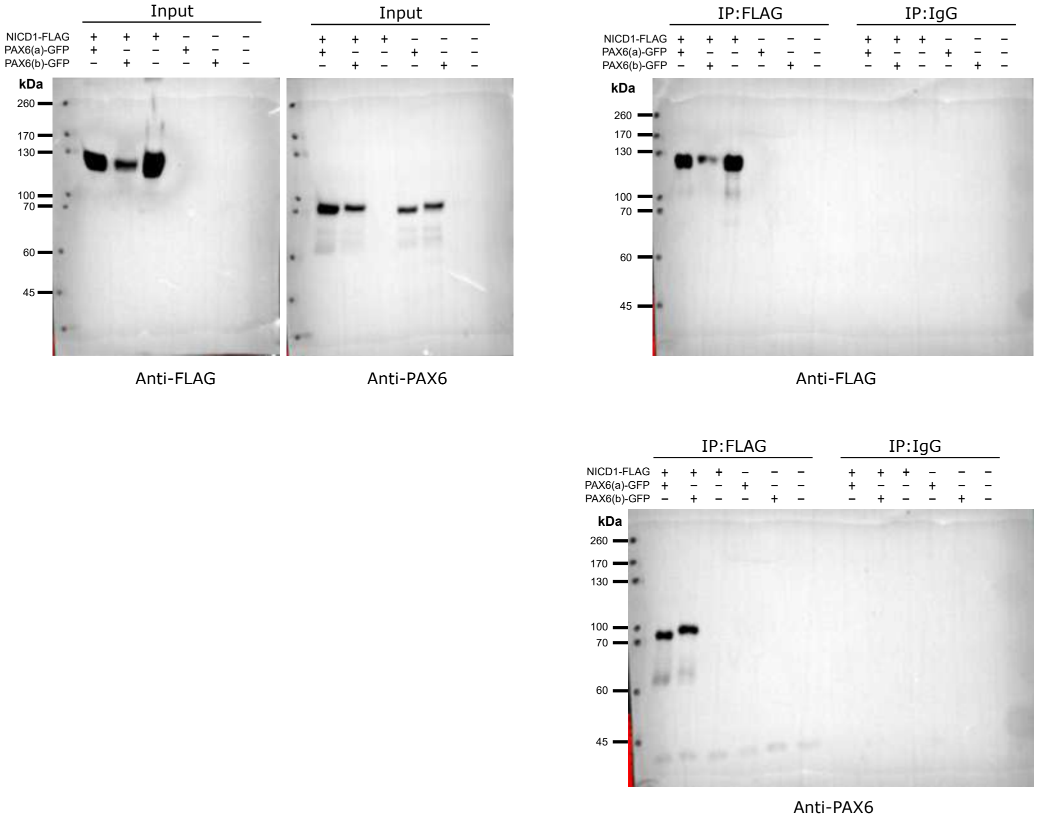
